# An Expanded Lpt Pathway Couples Surface Lipoprotein Export to Outer Membrane Vesicle Biogenesis in Bacteroides

**DOI:** 10.64898/2026.10.02.756314

**Authors:** Biswanath Jana, Evan J. Pardue, Tengfei Zhong, Mariana G. Sartorio, Manon Janet-Maitre, Juan C. Ortiz-Marquez, Marta Nieckarz, Tim Van Opijnen, Felipe Cava, Nichollas E. Scott, Mohammed Kaplan, Mario F. Feldman

## Abstract

The ecological dominance of Bacteroidota in the human gut microbiota derives from their exceptional capacity to metabolize diverse dietary and host-derived glycans. This ability relies on surface-exposed lipoproteins (SLPs), which are displayed on the bacterial surface and selectively packaged into outer membrane vesicles (OMVs). Although SLP export and OMV biogenesis are tightly coupled, the mechanism linking these processes has remained unknown. Here, we identify LptZ, a previously uncharacterized protein, as a critical link between SLP surface translocation and OMV biogenesis. Cryo-electron tomography of the *lptZ* mutant revealed chains of vesicles tethered to the outer membrane, offering a rare glimpse into an arrested stage of OMV formation in which vesicle fission is impaired. Proteomic analysis further showed that, in the absence of LptZ, a subset of SLPs fails to reach the cell surface and instead accumulates at the inner membrane. Unexpectedly, LptZ associates with the lipopolysaccharide transport (Lpt) machinery and repurposes it for SLP translocation. Crosslinking experiments revealed selective associations between multiple SLPs and Lpt proteins, providing mechanistic evidence for the direct involvement of the Lpt machinery in SLP trafficking. We further show that the *Bacteroides* Lpt machinery contains previously unrecognized components, revealing an unanticipated level of specialization relative to canonical Lpt systems. Together, our findings uncover a previously unknown protein export pathway, expand the functional role of the Lpt machinery, and establish a direct mechanistic link between SLP translocation and OMV biogenesis in Bacteroidota.

## Introduction

*Bacteroides* spp. are among the most abundant genera in the human gut microbiota, accounting for approximately 40% of bacterial species ^1–3^. Their ability to colonize the gut long-term is largely attributed to their capability to utilize a diverse range of complex carbohydrates from both dietary and host-derived sources ^4,5^. This carbohydrate utilization partly relies on the capacity of *Bacteroides spp*. to produce outer membrane vesicles (OMVs) ^6,7^. OMVs are spherical, membranous structures generated by the blebbing of the outer membrane (OM) ^8^. *Bacteroides fragilis* (*Bf*) and *Bacteroides thetaiotaomicron* (*Bt*) produce large quantities of uniformly sized OMVs, which carry enzymes essential for polysaccharide degradation ^6,9^. In other Gram-negative bacteria, OMVs play various roles in immune modulation, stress responses, and interbacterial interactions^8,10,11^. However, despite their physiological significance, the mechanisms underlying OMV biogenesis remain poorly understood.

Recent studies have begun to shed light on OMV biogenesis in Bacteroides ^12^. Proteomic analyses have revealed that OMVs are enriched in surface-exposed lipoproteins (SLPs), which primarily function as glycosyl hydrolases or proteases ^6,7,9^. Most of these lipoproteins contain a negatively-charged N-terminal motif, known as the Lipoprotein Export Signal (LES), which mediates both their surface exposure and packaging into OMVs ^9,13^. Furthermore, proteomic studies of *Bt* OMVs produced in the presence of different glycan nutrients have shown that *Bt* selectively modifies the composition of SLPs packaged into OMVs to optimize polysaccharide degradation ^7^. *Bacteroides spp* devote approximately 20% of their genomes to glycan breakdown and uptake, which are organized into distinct polysaccharide utilization loci (PULs) ^4,5^. PULs are clusters of colocalized and coregulated genes whose products coordinate the detection, enzymatic digestion, and transport of complex carbohydrates. Typical PULs contain an outer membrane β-barrel protein and one or more associated SLPs that facilitate the capture and degradation of polysaccharides before uptake by the cell ^14–16^.

Unlike *Bacteroides* spp, in species such as *Escherichia coli*, only a small subset of lipoproteins is surface-exposed. After translocation across the inner membrane (IM) via the Sec complex, lipoprotein transport is initiated by the lipidation of a conserved N-terminal cysteine residue to anchor the lipoprotein on the periplasmic side of the IM. Periplasmic lipoproteins then enter the Lol system and are translocated and inserted into the inner leaflet of the OM ^17,18^. However, the mechanisms by which SLPs cross the OM and reach their final destination on the cell surface remain unclear. In *Acinetobacter* and *Klebsiella*, a few SLPs have been shown to utilize the type 2 secretion system (T2SS) ^19,20^, while others, known as autotransporters, encode their own translocation machinery to cross the OM ^21–23^. In *Neisseria* and other bacterial species, a subset of SLPs is transported across the OM with the assistance of dedicated transporters called Surface Lipoprotein Assembly Modulators (Slam), which are typically encoded alongside their cognate SLPs ^24^. On the other hand, the OM of *Borrelia* species contains numerous SLPs, which have been proposed to reach the cell surface through the LPS transport (Lpt) machinery, a conserved multiprotein system that forms a continuous trans-envelope pathway and couples ATP hydrolysis at the inner membrane to the extraction, periplasmic transport, and insertion of LPS into the outer leaflet of the OM. Although *Borrelia* lacks LPS, it encodes several components of the Lpt system. In particular, the OM channel LptD has been proposed to mediate SLP translocation across the OM. However, this model remains controversial, as a separate study showed that LptD is dispensable for SLP transport^25,26^. A recent study demonstrated that crosstalk between the Lpt and Lol pathways is required for the surface localization of at least one lipoprotein, LptM, in *Pseudomonas aeruginosa* ^27^. Despite these insights, the export mechanisms have been elucidated for only a handful of SLPs, and a comprehensive, conserved pathway for SLP transport across bacterial species is yet to be established.

Through a high-throughput screen, we recently identified a novel family of anti-sigma factors that span both membranes and regulate OMV biogenesis in *Bt*^12^. The same screen also identified BT_3341, which we renamed LptZ (Lipoprotein transport protein Z), and found that LptZ is required for OMV production, prompting us to investigate its function. Furthermore, we show that LptZ is a component of an expanded Lpt machinery in *Bacteroides*, revealing an unexpected degree of specialization relative to canonical Lpt systems. Our findings demonstrate that this expanded Lpt machinery functions as an SLP export system, establishing a direct mechanistic link between SLP export and OMV biogenesis.

## Results

### LptZ is essential for OMV production

To identify proteins involved in OMV biogenesis, our lab previously developed a high-throughput transposon mutagenesis screen in *Bt*. This screen allowed us to identify several key genes critical for OMV production, including a mutant with a transposon insertion in gene *bt_3341*, hereafter referred to as *lptZ*, which appears to generate less OMV, compared to wild-type (WT)^12^. To investigate the role of LptZ in OMV biogenesis, we generated a clean deletion mutant of *lptZ* (referred to hereafter as ΔLptZ) and a corresponding complemented strain and analyzed the OMV production using SDS-PAGE (Fig. 1A). The ΔLptZ strain showed a significant decrease in both protein and LPS content within the OMV fraction compared to the WT strain (Fig. 1A-B). Moreover, the protein profile of the OMV fractions from the ΔLptZ differed substantially from that of the WT (Fig.1A). Comparative proteomic analysis confirmed these differences, as illustrated by principal component analysis (PCA), which showed a clear distinction between WT and ΔLptZ OMV profiles, while total membrane (TM) proteomes remained largely unchanged (Fig. 1C). Notably, the OMV fraction from ΔLptZ lacked the canonical OMV-associated proteins and was instead enriched in cytoplasmic proteins, including ribosomal components, suggesting that most of these are lysis-derived vesicles (LV), originating from cell lysis (Fig. 1D; Table S1; Dataset 1). Constitutive expression of LptZ in the mutant background restored both OMV production and composition to WT levels (Fig. 1A-C). Collectively, these findings demonstrate that LptZ is essential for OMV biogenesis. Although we cannot exclude that a small number of bona fide OMVs are produced by the ΔLptZ strain, the vast majority of vesicular material recovered from the OMV fraction appears to originate from cell lysis rather than from the canonical OMV biogenesis pathway. Thus, deletion of *lptZ* reduces bona fide OMV production to near-undetectable levels. To our knowledge, this represents the first reported bacterial strain with such a profound defect in OMV biogenesis.

**Figure 1:**
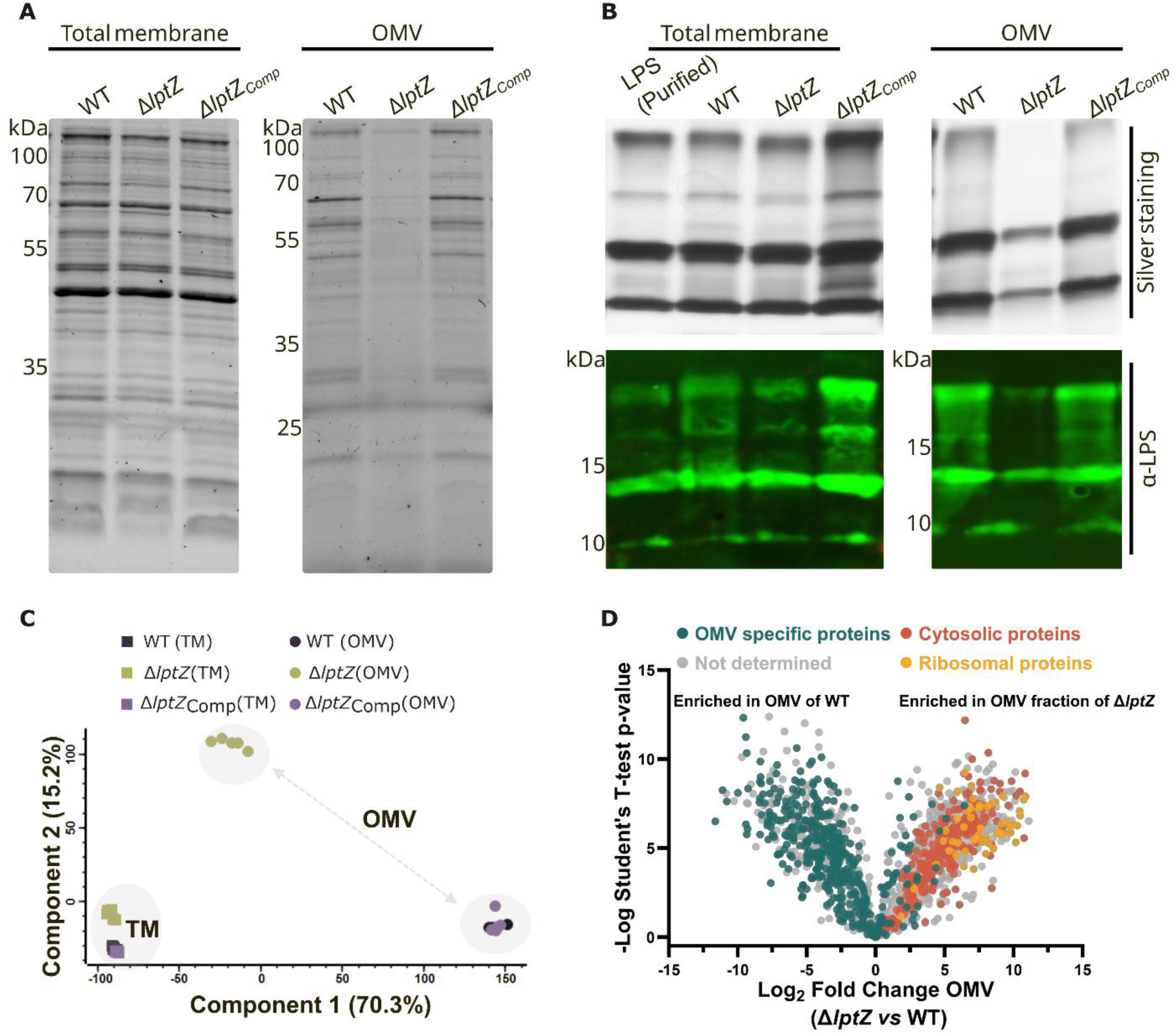
LptZ is essential for OMV secretion in *Bt*. **(A)** SDS-PAGE analysis of total membrane (TM) and outer membrane vesicle (OMV) protein content. TM and OMVs were purified from WT, Δ*lptZ* mutant, and complemented (Δ*lptZ*comp) strains and subjected to SDS-PAGE analysis. Protein samples were normalized by OD_600_ and loaded onto a 12% SDS-PAGE gel. **(B)** LPS analysis of TM and OMV fractions. LPS was isolated from all three strains and analyzed by 10–20% SDS-PAGE, followed by silver staining and Western blot analysis using anti-LPS antibodies. Experiments were independently repeated at least three times with similar results; one representative result is shown. **(C)** Principal Component Analysis (PCA) of TM and OMV proteomes. TM and OMVs were purified from all three strains and subjected to proteomic analysis; the data were analyzed using Perseus software. **(D)** Comparison of OMV proteomes. The Δ*lptZ* OMV fraction lacked OMV-specific proteins (teal circles) and showed a significant enrichment of cytosolic proteins (orange circles), including ribosomal proteins (amber circles), indicating cell lysis rather than regulated OMV release in the mutant.

### Loss of LptZ leads to stalled OMV biogenesis at the membrane scission step

To further investigate the role of LptZ in OMV biogenesis, we examined WT and ΔLptZ cells by cryo-electron tomography (cryo-ET), which enables visualization of cellular structures in their native, hydrated state at nanometer resolution^28^. WT cells exhibited a consistent rod-shaped morphology, with a well-defined periplasmic space and intact OM, and produced OMVs of relatively uniform size (Fig. 2A-C and Fig. S1A-B). In contrast, ΔLptZ cells displayed pronounced morphological abnormalities, including expansion of the periplasm, a more rounded cell shape, and elongated tubular structures extending from the OM (Fig. 2D-G and Fig. S1C-I). Strikingly, we also observed chains of vesicle-like structures that remained tethered to the OM and appeared as contiguous extensions of the outer envelope (Fig. 2E-F). These observations indicate that early stages of OMV biogenesis, including OM curvature and budding, can occur in the absence of LptZ, whereas the subsequent membrane scission step is severely impaired. The accumulation of these stalled intermediates provides morphological evidence of an otherwise transient stage of OMV biogenesis. Together, these findings indicate that the release of nascent vesicles from the bacterial surface requires LptZ.

**Figure 2:**
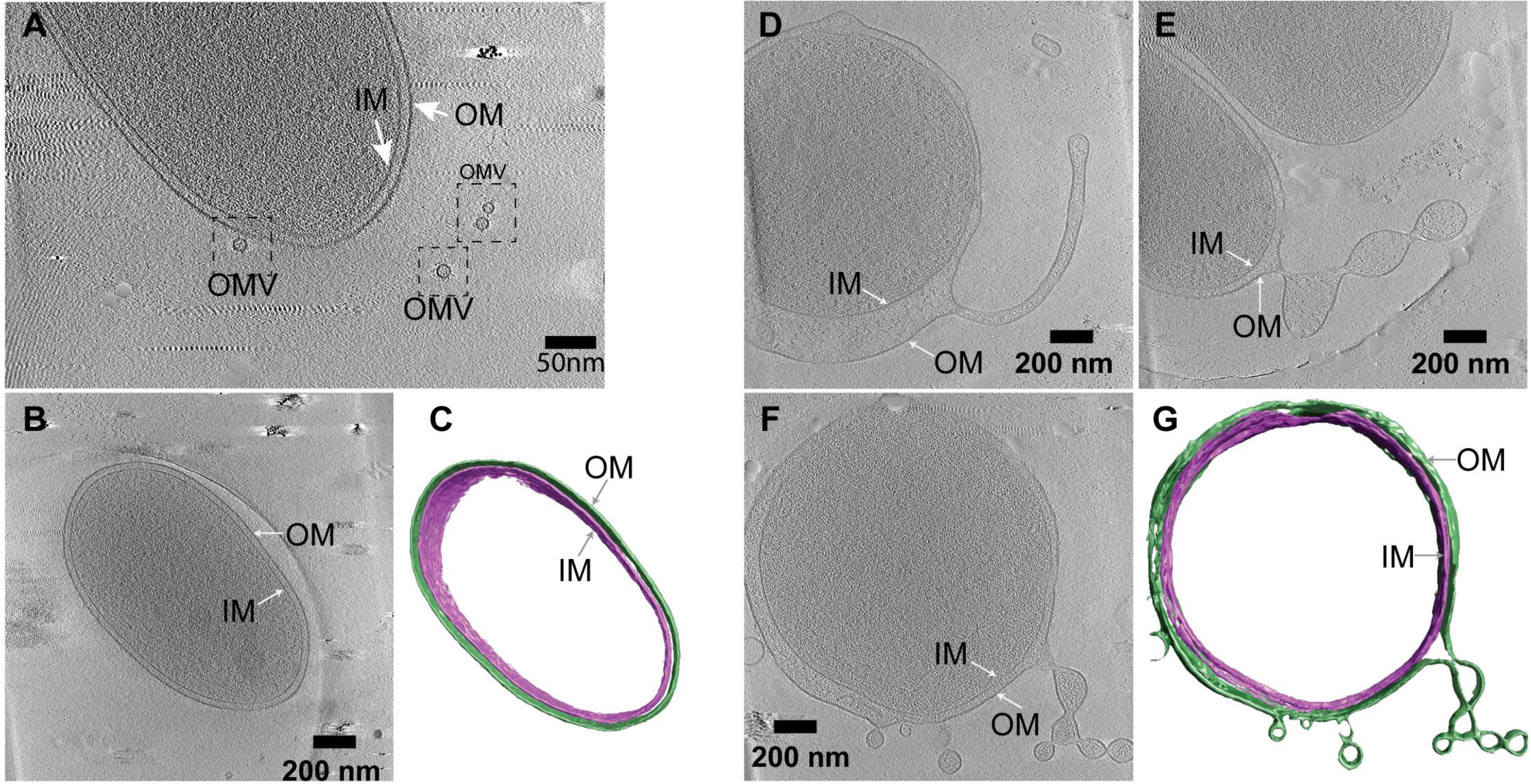
Loss of LptZ impairs OMV formation. (**A-B)** A slice through cryo-electron tomogram of a WT *Bt* cell showing the bacilli shape of the cell and OMVs secreted from the cell. (**C)** 3D segmentation of the cell shown in (B) with the OM shown in green and IM in purple. Scale bar is 200 nm. (**D-F)** Similar to (A-B) but for Δ*lptZ* mutant cells, illustrating the irregular shape of the cells and the presence of vesicle-like structures attached to the OM. Scale bar is 200 nm. (**G)** 3D segmentation of the cell shown in (F) with the OM shown in green and the IM in purple. Scale bar is 200 nm.

### LptZ is a component of an expanded Lpt machinery

LptZ is a 642-amino-acid protein. AlphaFold predicts that LptZ comprises five distinct domains, preceded by an N-terminal signal peptide. The N-terminal region, domain 1, appears to adopt an immunoglobulin-like (Ig-like) fold, while the C-terminal domain (domain 5) aligns with the DUF2141 (Fig. 3A). Notably, none of the five domains has an assigned or predicted function, suggesting that LptZ represents a previously uncharacterized protein architecture. Previous reports revealed that LptZ is essential for *Bt* survival in the murine gut ^29,30^. However, the molecular mechanism underlying this requirement remains unclear. To elucidate the precise subcellular localization of LptZ, we first isolated IM and OM fractions from *Bt* and performed comparative proteomic analysis. The data revealed that LptZ is present predominantly in the IM fraction (Fig. 3B; Dataset 2). We further validated this result by expressing 3xFLAG-tagged LptZ in *Bt* and isolating total membranes (inner and outer; TM), IM, and OM fractions prior to Western blotting. SecA (BT_4362) and SusC were used as IM and OM controls, respectively. FLAG-tagged LptZ was most abundant in the IM fraction (Fig. S2).

**Figure 3:**
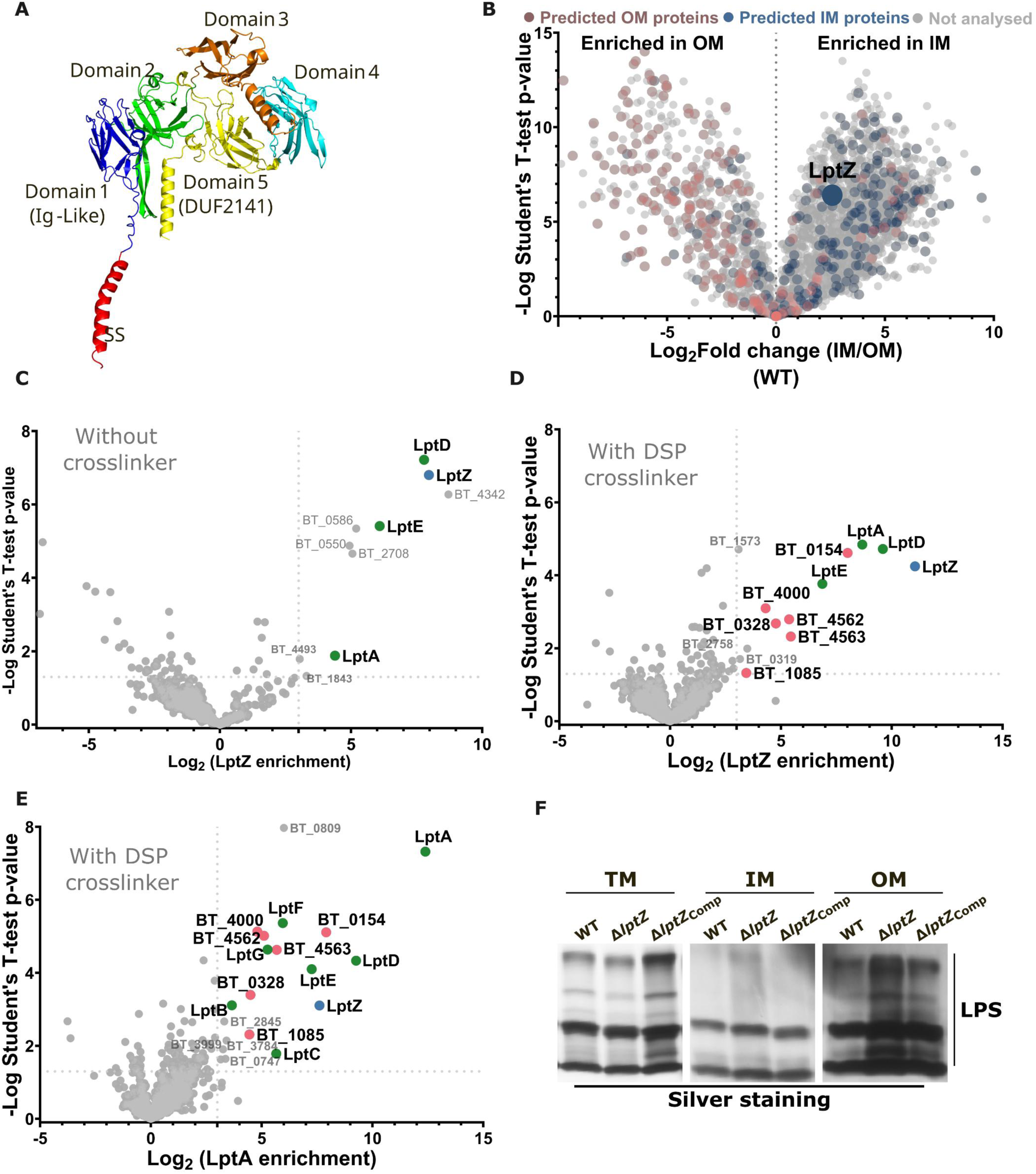
LptZ is an IM protein and associates with the Lpt machinery but is not required for LPS transport. **(A)** AlphaFold-predicted structure of LptZ showing an N-terminal signal peptide (red) and five domains: Domain 1 (blue) adopts an Ig-like fold; Domains 2 (green), 3 (bronze), and 4 (cyan); Domain 5 (yellow) aligns with DUF2141. **(B)** Volcano plot of isolated IM and OM proteomes of *Bt* (WT), highlighting enrichment of predicted IM-specific proteins (blue circles) and OM-specific proteins (maroon circles) in their respective membrane fractions. LptZ (blue circle) is mostly detected in the IM fraction. **(C)** Volcano plot showing proteins enriched during affinity purification of LptZ from *Bt* (without crosslinker) expressing a C-terminal dual-tagged (3xFLAG-His₁₀-tagged) LptZ construct, followed by sequential Ni-NTA and anti-FLAG affinity purification. **(D, E)** Volcano plots showing proteins enriched during affinity purification of LptZ (D) or LptA (E) following *in vivo* crosslinking with DSP. Both LptZ and LptA were C-terminally 3xFLAG-tagged and purified from membrane using anti-FLAG affinity resin. Mass spectrometry identified several proteins significantly enriched with LptZ or LptA relative to the corresponding WT controls, suggesting potential LptZ or LptA-interacting partners. **(F)** LPS from TM, IM, and OM fractions of WT, Δ*lptZ* mutant, and complemented *Bt* strains was isolated and separated by 10-20% SDS-PAGE and visualized by silver staining. Experiments were independently repeated at least three times with similar results; one representative result is shown.

To investigate the potential role of LptZ in OMV biogenesis, we sought to identify its interacting partners. We first expressed a genomically integrated, C-terminally 3xFLAG-His10-tagged LptZ from a constitutive promoter in the Δ*lptZ* background. Tagged LptZ was functional, restoring OMV production to WT levels (Fig.S3). Membrane-associated LptZ was sequentially purified using Ni-NTA and anti-FLAG affinity chromatography, followed by mass spectrometry to identify co-purifying proteins (Fig. 3C; Dataset 3). To preserve higher-order complexes and capture transient or weak interactions, we also generated a genomically integrated 2xFLAG-tagged LptZ strain and performed *in vivo* DSP crosslinking before anti-FLAG affinity purification and mass spectrometry (Fig. 3D; Table S2; Dataset 4). An isogenic WT strain lacking tagged LptZ served as the negative control in both experiments. Three known components of the LPS transport machinery, LptD (BT_2133), LptE (BT_4376), and LptA (BT_3849), co-purified with LptZ in both crosslinked and non-crosslinked experiments (Fig. 3C-D, Table S2, Dataset 3-4)^31^. Notably, LptD and LptA were also predicted as LptZ interactors in our STRING network analysis (Fig. S4), providing independent support for these interactions. Together, these findings indicate that LptZ associates with the LPS transport machinery.

Because affinity purification of LptZ from *Bt* could capture indirect associations mediated by other components of the Lpt machinery, we next sought to identify which Lpt proteins interact directly with LptZ. We therefore tested LptZ against all seven known Lpt components, LptA, LptB, LptC, LptD, LptE, LptF, and LptG, using heterologous expression in *E. coli*. His-tagged versions of each Lpt protein were co-expressed with or without LptZ-FLAG, and anti-FLAG pull-down assays were performed to assess direct interactions. This approach confirmed direct interactions of LptZ with LptD and LptE and additionally revealed interactions with LptF and LptG (Fig. S5A-D and 5H). In contrast, LptA, LptB, and LptC were not detected as interactors in this assay (Fig. S5E-H). These interactions were further evaluated using bacterial adenylate cyclase two-hybrid (BACTH) assays (Fig. S6). LptB and LptC produced positive signals only in the BACTH assay, whereas LptD, LptE, LptF, and LptG were identified by both approaches (Fig. S5-6). The concordant detection of LptD, LptE, LptF and LptG by both Co-IP and BACTH provided the strongest evidence for their direct interaction with LptZ. These results establish an association between LptZ and the core Lpt components.

We further tested the association between LptZ and the Lpt machinery by purifying LptA, a central component of the Lpt complex. A genomically integrated 2xFLAG-tagged LptA was generated in WT *Bt*, followed by *in vivo* DSP crosslinking, anti-FLAG affinity purification, and mass spectrometry. Major components of the Lpt machinery, including LptB, LptC, LptD, LptE, LptF, and LptG, co-purified with LptA, validating the effectiveness of this approach (Fig. 3E; Table S2; Dataset 5). Notably, LptZ was also significantly enriched in the LptA pull-down, confirming independently that LptZ associates with the Lpt machinery (Fig. 3E; Table S2; Dataset 5). In addition to the known Lpt components, six previously uncharacterized proteins-BT_0154, BT_4000, BT_4562, BT_4563, BT_0328, and BT_1085-co-purified with both LptZ and LptA under crosslinking conditions (Table S2; Dataset 4 and 5). BT_0154 is predicted to encode a periplasmic protease, whereas the remaining proteins lack assigned functions or recognizable functional domains. Importantly, the co-purification of these proteins with both LptZ and LptA suggests that they may represent previously uncharacterized components or associated factors of an expanded Lpt machinery. Importantly, despite its association with the Lpt machinery, deletion of *lptZ* does not impair LPS transport to the OM or alter LPS size or profile (Fig. 3F). This unexpected finding indicates that LptZ is not required for the canonical LPS transport function of the Lpt system and instead suggests that it contributes to a distinct activity of the machinery.

### LptZ is necessary for surface localization of Sus proteins and growth in complex glycans

Given its association with the Lpt machinery and requirement for OMV biogenesis, we next examined whether loss of LptZ affects OM integrity. We tested the ΔLptZ mutant against several OM-disrupting agents, including EDTA, vancomycin, and polymyxin B. Although the mutant grew normally on brain heart infusion (BHI) agar under non-stress conditions (Fig. S7), growth was markedly impaired in the presence of all three OM-stressor compounds. These findings indicate that loss of LptZ compromises OM integrity and increases susceptibility to envelope stress, supporting a critical role for LptZ in OM homeostasis.

Bacteroides species encode a large repertoire of SLPs that contribute to polysaccharide degradation and growth on complex glycans, and these proteins are highly enriched in OMVs^6,7,9^. Because LptZ is required for OMV biogenesis yet is dispensable for LPS transport, we reasoned that its association with the Lpt machinery might confer an additional function on this complex. The requirement for LptZ in both SLP-enriched OMV secretion and maintenance of OM integrity suggested that LptZ may associate with the Lpt machinery for SLP trafficking, facilitating their transport to the cell surface and subsequent incorporation into OMVs. To test this hypothesis, we first assessed whether loss of LptZ compromises SLP-dependent utilization of complex glycans. ΔLptZ exhibited growth comparable to WT in rich medium and only a prolonged lag phase on simple sugars such as glucose and maltose (Fig. 4A). In contrast, ΔLptZ failed to grow on complex polysaccharides, including starch, heparin, and hyaluronan (Fig. 4A), consistent with a defect in the surface localization of the SLPs required for their degradation. To further investigate the defects in polysaccharide utilization and SLP expression observed in the ΔLptZ mutant, we examined the well-characterized sus operon, which encodes the known SLPs involved in starch binding and utilization: SusD (BT_3701), SusE (BT_3700), SusF (BT_3699), and SusG (BT_3698) ^14,16^. The protein abundance of all four Sus SLPs were markedly reduced in ΔLptZ, even under starch-inducing conditions (Fig. 4B). Complementation of the ΔLptZ restored the levels of these proteins to those observed in the WT strain (Fig. 4B). Interestingly, transcriptomic analysis revealed no significant differences in transcription of the Sus genes between the WT and ΔLptZ (Fig. S8; Dataset 6). These findings suggest that the Sus SLPs (and possibly other SLPs as well) are expressed but may fail to reach their proper location, indicating that defective SLP trafficking results in mislocalization and subsequent degradation.

**Figure 4:**
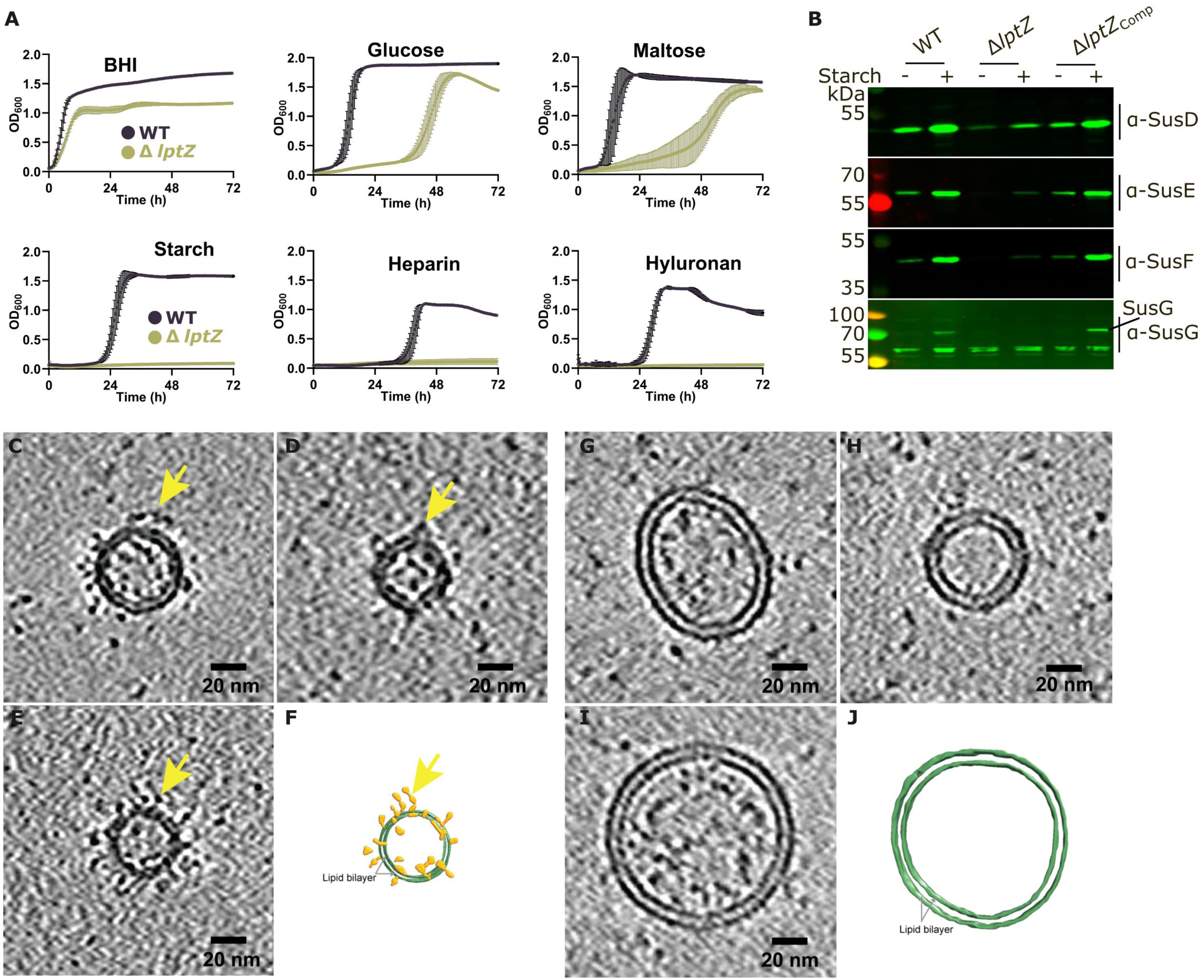
LptZ is required for the trafficking of SLPs to the cell surface. **(A)** Growth curves of *Bt* WT (purple) and Δ*lptZ* (yellow-green) in minimal media containing the indicated carbon sources or in rich BHI medium. **(B)** Immunoblot analysis of total protein lysates from *Bt* WT, Δ*lptZ*, and complemented strains cultured in BHI with or without 0.5% (w/v) starch. Blots were probed with custom polyclonal antibodies against SusD, SusE, SusF, and SusG. Starch supplementation robustly induced Sus proteins in the WT strain, whereas their expression was markedly reduced in the Δ*lptZ* mutant under the same conditions. SusG-specific band is indicated by lines and labeled as “SusG.”. Experiments were independently repeated at least three times with similar results; one representative result is shown.**(C-J)** Cryo-EM of OMVs and lysis-derived vesicles (LVs) from *Bt* strains. (C-E) Slice through cryo-electron tomograms of OMVs derived from WT *Bt* cells (from Figure 2A), highlighting the presence of densities associated with these OMVs (yellow arrows). Scale bar is 20 nm. (F) 3D segmentation of the OMV displayed in (E) with the membranes shown in green and associated densities shown in yellow. (**G-J)** Similar to panels C-F, but for the LVs isolated from the Δ*lptZ* mutant. Scale bar is 20 nm.

Cryo-ET analysis of OMVs isolated from WT bacteria revealed abundant surface-associated densities, most of which projected outward from the vesicle membrane (Fig. 4C-F). Given that *Bt* OMVs are highly enriched in SLPs^6,9^, these densities are consistent with surface-exposed SLPs. In striking contrast, cryo-ET analysis of LVs isolated from ΔLptZ cells revealed lipid bilayers that were largely devoid of the surface densities observed on WT OMVs (Fig. 4G-J). The absence of these structures from ΔLptZ vesicles further supports their assignment as SLPs and is consistent with a defect in SLP surface localization. LVs were also morphologically distinct from WT OMVs: whereas WT OMVs exhibited a relatively uniform size distribution, ΔLptZ LVs were significantly larger and displayed substantially greater size heterogeneity (Fig. S9; Dataset 7). Together, these size differences further distinguish the LVs recovered from ΔLptZ cells from bona fide OMVs isolated from WT cells and further confirm our previous findings that these LVs originate from cell lysis.

### LptZ is essential for the surface localization of a subset of SLPs

To determine the extent to which LptZ influences SLP trafficking, we isolated IM, OM, and OMV fractions from WT and ΔLptZ cells and performed comparative proteomic analysis. As in WT cells, fractionation of ΔLptZ cells yielded well-separated IM and OM fractions, with most IM and OM marker proteins enriched in their respective fractions (Fig. S10; Dataset 2). We integrated these data with RNA sequencing to distinguish changes in protein localization from changes in gene expression. Because SusD-like proteins represent a major class of Bacteroides SLPs with established surface localization^13,16^, we first used them as a proxy for SLP trafficking. Of 97 SusD-like proteins, 66 were detected by proteomics and included in the analysis. Although transcript levels of most SusD-like homologs remained largely unchanged in ΔLptZ (RNA fold changes between -1 and +1), their abundance in the OM was markedly reduced (Fig. 5A; Dataset 8). Consistent with a trafficking defect, most SusD-like proteins showed an increased IM-to-OM ratio in ΔLptZ compared with WT, indicating that their delivery to the OM is affected in the absence of LptZ, resulting in their accumulation in the IM (Fig.5B, Dataset 8). A smaller subset showed unchanged IM-to-OM ratios in the mutant, suggesting that their trafficking is LptZ-independent. Collectively, these findings indicate that LptZ is required for efficient translocation of most SusD-like SLPs from the IM to the OM (Fig. S11A).

**Figure 5:**
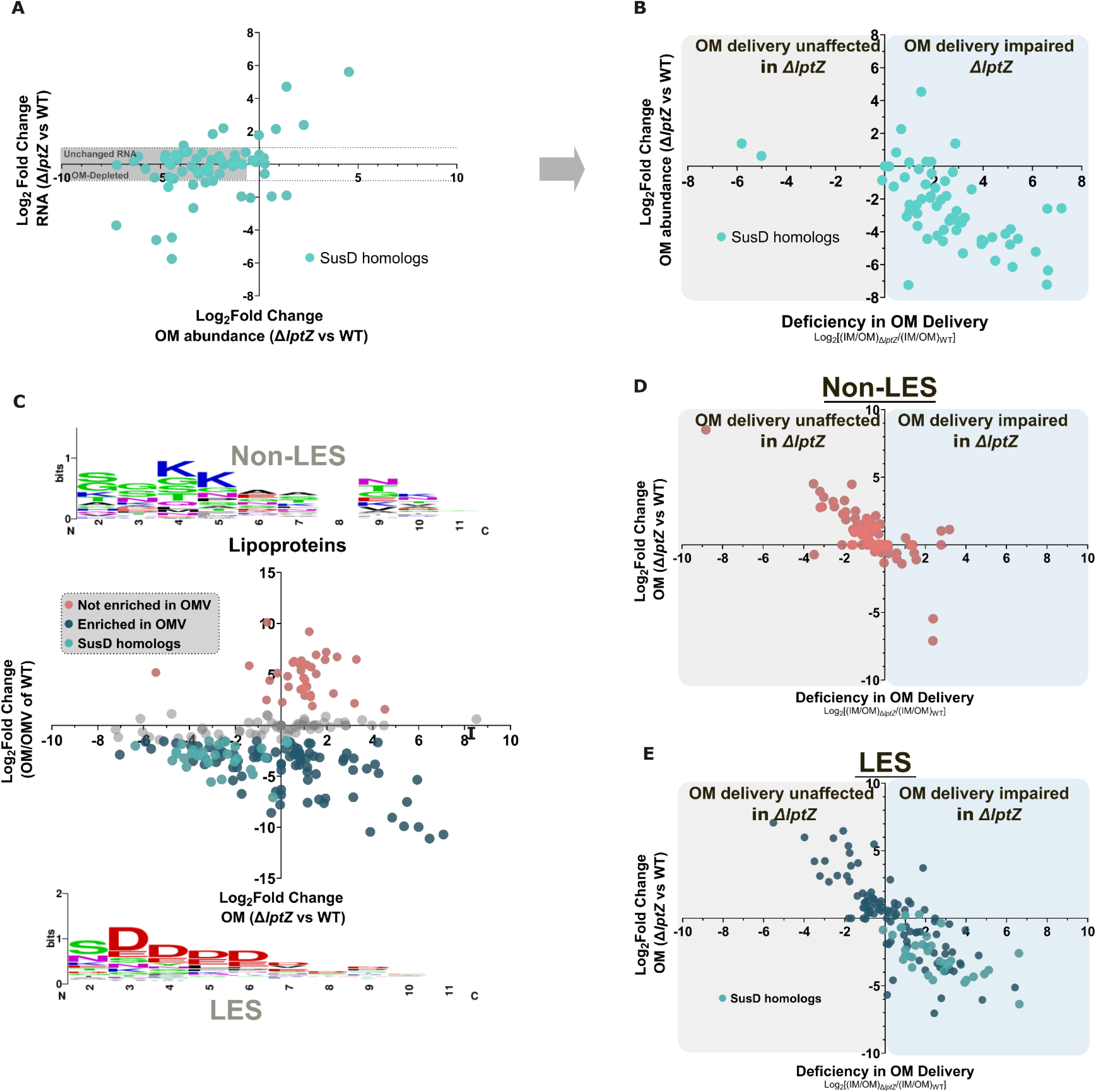
LptZ is essential for the surface localization of a subset of SLPs. **(A)** Integrated transcriptomic and OM proteomic analysis of SusD-like lipoproteins (cyan circles) between the Δ*lptZ* mutant and WT. Y-axis shows log₂ fold change in transcript levels; the x-axis shows log₂ fold change in OM protein abundance. Dotted lines (±1 log₂ fold) denote thresholds for unchanged transcript expression between the Δ*lptZ* mutant and WT. Most SusD-like proteins display unchanged transcript levels but significantly reduced OM abundance in the Δ*lptZ* mutant, indicating post-transcriptional mislocalization. **(B)** To assess trafficking defects of SusD-like lipoproteins, the relative change in subcellular distribution between IM and OM was calculated as log₂[(IM/OM ratio in Δ*lptZ*) / (IM/OM ratio in WT)] and plotted against the OM abundance. Positive values on the x-axis indicate increased IM retention and impaired OM delivery in the Δ*lptZ* mutant, while values near zero or negative reflect unaffected localization. Most SusD-like lipoproteins exhibit elevated IM/OM ratios and lower OM abundance in Δ*lptZ*, confirming LptZ-dependent trafficking. **(C)** Lipoproteins with unchanged transcript levels were selected for this analysis. OMV-enriched lipoproteins (deep teal circles) and OM-retained lipoproteins (soft red circles) were classified based on their relative enrichment in OM versus OMVs (y-axis) and plotted against OM abundance (x-axis). Lipoproteins not strongly enriched in either OM or OMVs are shown as grey circles. Sequence logos of the ten residues immediately downstream of the conserved lipoprotein attachment cysteine reveal a conserved LES motif in OMV-enriched proteins (bottom), and a distinct Non-LES motif in OM-retained proteins (top). **(D-E)** Lipoproteins were classified based on the presence of either a Non-LES (panel D) or LES (panel E) motif. Trafficking defects in each lipoprotein subset were assessed by plotting log₂[(IM/OM ratio in Δ*lptZ*) / (IM/OM ratio in WT)] (x-axis) against OM abundance (y-axis). (D) Non-LES lipoproteins (soft red), which are typically OM-retained, show unchanged or reduced IM/OM ratios in Δ*lptZ*, and stable OM abundance, indicating LptZ-independent localization. (E) A subset of LES-containing lipoproteins (deep teal), including SusD homologs (cyan), exhibit increased IM/OM ratios and reduced OM abundance in Δ*lptZ*, consistent with LptZ-dependent trafficking. In contrast, other LES lipoproteins maintain lower or unchanged IM/OM ratios and stable OM abundance, suggesting LptZ-independent trafficking. Figure S11 shows a schematic overview of the data.

To determine whether the observed localization defect is specific to SusD-like SLPs or reflects a broader disruption of lipoprotein trafficking, we extended our analysis to all previously annotated lipoproteins, including SusD-like proteins^32^. We focused on lipoproteins whose transcript levels were unchanged in ΔLptZ (log_2_ fold change between -1 and +1), thereby minimizing the contribution of altered gene expression to differences in protein localization. We first examined lipoprotein partitioning in WT cells to identify proteins preferentially enriched in OMVs versus retained in the OM (Fig. 5C; Dataset 9). Sequence logos generated from the ten residues immediately following the lipidated cysteine revealed, as expected, that OMV-enriched proteins (Fig. 5C, deep teal blue circles) contained the LES motif, whereas this motif was absent from OM-retained lipoproteins (Fig. 5C, soft red circles) ^9,13^.

We next examined the effect of LptZ deletion on these two classes of lipoproteins. Most non-LES lipoproteins retained similar IM-to-OM distributions in ΔLptZ and WT cells, indicating that their trafficking to the OM is largely LptZ-independent (Fig. 5D; Fig. S11B, Dataset 10). In contrast, most LES-containing lipoproteins showed increased IM-to-OM ratios in ΔLptZ, consistent with defective translocation from the IM to the OM (Fig. 5E, Fig. S11C, Dataset 11). A smaller group of LES-containing lipoproteins maintained WT-like OM abundance and IM-to-OM ratios in the absence of LptZ, indicating that not all SLPs require LptZ for trafficking. Together, these proteomic analyses indicate that LptZ is required for the efficient export of most SLPs, whereas trafficking of non-LES lipoproteins is largely unaffected by its absence. This conclusion is consistent with recent work by Armbruster et al.^33^, showing that SLP export does not depend on a functional Lol system. Consistent with these findings, LptZ disruption and LolB depletion affected largely distinct sets of lipoproteins (Fig. S12A-B; Dataset 12). Notably, most SusD proteins were strongly affected by LptZ deletion but showed little change upon LolB depletion (Fig. S12C; Dataset 13). These results indicate that LES-containing SLPs and non-LES lipoproteins have distinct trafficking requirements, with LptZ specifically required for the efficient export of most LES-containing SLPs.

### Lpt machinery mediates SLP transport

The findings described in the previous sections raised the possibility that LES-containing SLPs are transported by an Lpt machinery-dependent pathway. To test this hypothesis, we performed *in vivo* crosslinking followed by affinity purification. We examined two classes of lipoproteins: the LES-containing SusD and SusF proteins encoded by the maltose-inducible BT_3698-BT_3705 locus^14^, and three non-LES lipoproteins, BT_0705, BT_2548, and BT_4399. These non-LES lipoproteins localize to the OM but are not predicted to be surface exposed, and their trafficking to the OM is LptZ-independent (Dataset 10). For these experiments, LptA and LptZ were chromosomally tagged with C-terminal 2xFLAG and used as bait proteins, whereas the three non-LES lipoproteins were chromosomally tagged with C-terminal His_10_ expressed from a constitutive promoter^34^. SusD and SusF remained untagged and were detected using specific antibodies. Following *in vivo* crosslinking with DSP and anti-FLAG affinity purification, SusD and SusF were detected in complexes containing both LptA and LptZ. In contrast, none of the three His-tagged non-LES lipoproteins were detected in LptA- or LptZ-containing complexes (Fig. 6A; Fig.S13A-B). These findings indicate that LES-containing lipoproteins, but not non-LES lipoproteins, specifically engage the Lpt machinery during trafficking.

**Figure 6:**
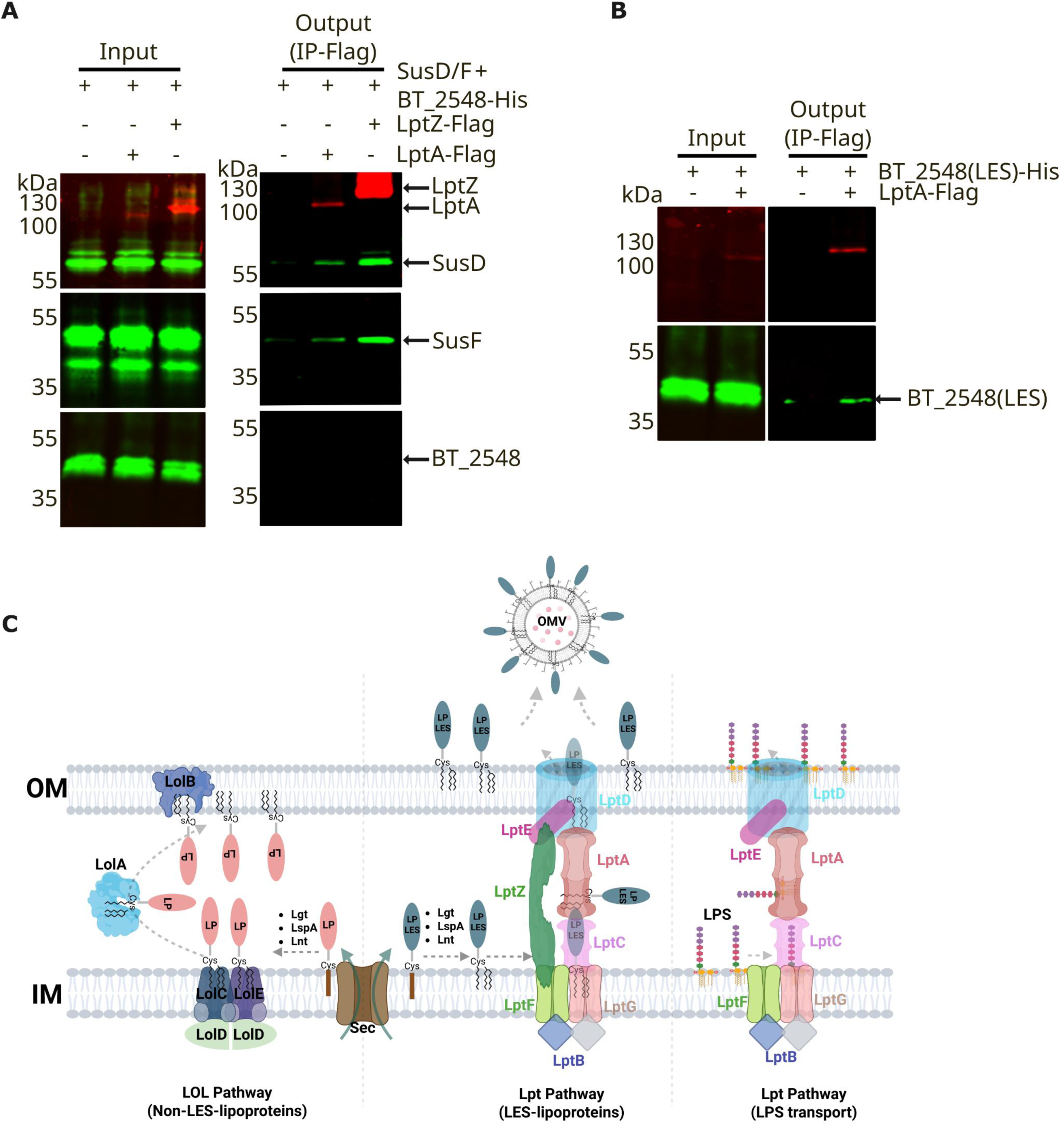
LptZ promotes SLP trafficking through the Lpt machinery. **(A)** *In vivo* crosslinking with DSP followed by co-purification of LptA and LptZ was used to check the association of LES-containing and non-LES-containing lipoproteins with the Lpt machinery. LptA and LptZ were genomically tagged at their C-termini with 2xFLAG and used as bait proteins. The LES-containing lipoproteins SusD and SusF were induced with maltose, whereas the non-LES outer-membrane lipoprotein BT_2548 was genomically expressed with a C-terminal His-tag from a constitutive promoter. SusD and SusF were detected using specific antibodies, and BT_2548 was detected using an anti-His antibody. Following DSP crosslinking, SusD and SusF co-purified with both LptA and LptZ, relative to the untagged WT control, whereas BT_2548 was not detected with either LptA or LptZ. **(B)** The native non-LES domain of BT_2548 was replaced with an LES domain to generate BT_2548(LES). In contrast to BT_2548, BT_2548(LES) co-purified with LptA, demonstrating that incorporation of the LES domain is sufficient to promote association of BT_2548 with the Lpt machinery. Experiments were independently repeated at least three times with similar results; one representative result is shown. **(C)** Proposed lipoprotein transport mechanism in the Bacteroidota phylum. Lipoproteins (LP) are first translocated across the IM and undergo enzymatic maturation, including signal peptide cleavage and acylation. Lipoproteins that remain anchored to the inner leaflet of the OM (Non-LES, soft red ellipse) are delivered via the canonical Lol pathway. In contrast, SLPs (deep teal ellipse) containing a conserved LES motif are trafficked by the Lpt machinery in complex with LptZ and subsequently incorporated into OMVs. Hence, LptZ co-opts the Lpt complex to transport SLPs directly from the IM to the cell surface, similar to LPS translocation. The core Lpt machinery, without LptZ, is responsible for LPS transport. This model illustrates the dual functionality of the Lpt complex, governed by the presence or absence of LptZ.

To test whether the LES sequence is sufficient to redirect a non-LES lipoprotein to the Lpt pathway, we replaced the native N-terminal region of BT_2548, including its signal peptide and non-LES sequence, with the corresponding signal peptide and LES-containing region of SusD. Introduction of the SusD LES sequence substantially increased the abundance of BT_2548 in the OMV fraction without altering overall OMV production (Fig. S14). Moreover, LES-containing BT_2548 co-purified with LptA (Fig. 6B), demonstrating that acquisition of the LES sequence is sufficient to redirect BT_2548 toward the Lpt-dependent trafficking pathway.

Together, these findings support a model in which Bacteroides uses distinct pathways to traffic different classes of lipoproteins: the classical Lol pathway primarily delivers non-LES lipoproteins to the OM, whereas the Lpt machinery, including LptZ, selectively engages LES-containing lipoproteins and mediates their translocation to the cell surface and subsequent enrichment in OMVs (Fig. 6C).

### LptZ is a Bacteroidota-specific component of the Lpt machinery

BLAST analyses revealed that LptZ homologs are broadly conserved across the phylum Bacteroidota, with representatives in the orders Bacteroidales, Cytophagales, Flavobacteriales, Marinilabiliales, and Sphingobacteriales (Fig. S15A-B, Dataset 14). In contrast, LptZ homologs were not detected in other major Gram-negative lineages, including Proteobacteria, suggesting that LptZ represents a Bacteroidota-specific adaptation of the canonical Lpt machinery associated with the distinctive cell-surface biology of this phylum. Thus, Bacteroidota appear to have expanded the function of an ancient LPS trans-envelope transport system to transport an entirely different class of lipidated proteins.

## DISCUSSION

Despite decades of study, the mechanisms governing OMV biogenesis in Gram-negative bacteria remain poorly defined. Although several mechanisms have been proposed, many are based on mutants with altered envelope homeostasis that exhibit increased vesiculation. Here, we identify LptZ as a Bacteroidota-conserved protein required for OMV production in *B. thetaiotaomicron*. Deletion of *lptZ* nearly abolished bona fide OMV secretion. The vesicular material recovered from ΔLptZ cultures lacks the SLPs that constitute the canonical cargo of Bacteroides OMVs and instead contains ribosomal and cytoplasmic components, indicative of cell lysis (Fig. 1). Strikingly, cryo-ET revealed chains of interconnected, incompletely separated vesicles that remained associated with the ΔLptZ cell surface (Fig. 2D-G). These structures support a two-step model of OMV biogenesis in which membrane curvature and budding occur first, followed by membrane scission and vesicle release. Whereas the initial budding step can proceed in the absence of LptZ, the subsequent scission and release step is severely impaired. We further show that LptZ is a component of an expanded Lpt transport system. LptZ is dispensable for LPS export but required for the efficient trafficking of most LES-containing SLPs. Together, these findings reveal an unexpected connection between SLP trafficking and OMV biogenesis and support a model in which Bacteroidota have expanded the function of the canonical Lpt machinery to accommodate the transport of a large repertoire of SLPs (Fig 6C).

Lipoproteins are matured at the IM and, in most Gram-negative bacteria, are transported to the OM by the well-characterized Lol system, which delivers them to the periplasmic leaflet of the OM. Two recent studies support the existence of an alternative pathway for SLP trafficking in Bacteroidota that does not require the canonical LolA/LolB route^33,35^. In *Flavobacterium johnsoniae*, loss of LolA or LolB does not abolish SLP surface localization^35^. Similarly, depletion of LolB in *B. thetaiotaomicron* does not substantially affect SLP trafficking^33^ (Fig.12). Our results identify an expanded Lpt machinery as a pathway responsible for SLP surface trafficking in *B. thetaiotaomicron*. This model is conceptually related to observations in *Borrelia*, where components of the Lpt machinery have been proposed to participate in SLP surface localization despite the absence of LPS^26^. Unlike *Borrelia*, however, *Bacteroides* must transport both LPS and a large repertoire of SLPs, suggesting that incorporation of LptZ may have broadened the function of the canonical Lpt machinery to mediate SLP transport.

Most SLPs fail to reach the cell surface in the absence of LptZ and are retained in the IM instead. Consistent with this trafficking defect, our crosslinking experiments detected the LES-containing SLPs including SusD and SusF in association with components of the Lpt machinery. We propose that these interactions capture SLPs during their transit from the IM to the OM. In contrast, the non-LES lipoproteins that were co-expressed and purified from the same cells as SLPs examined in this study were not detected in association with the Lpt machinery (Fig 6A; Fig S13). Moreover, replacing the N-terminal region of a non-LES lipoprotein with the LES-containing N-terminal region of SusD promoted its association with the Lpt machinery and enhanced its incorporation into OMVs, consistent with redirection toward the Lpt-dependent trafficking pathway (Fig 6B; Fig S14). Together, these findings indicate that the information contained within the N-terminal region of SLPs determines their entry into the LptZ-dependent trafficking pathway.

Our crosslinking experiments identified several additional proteins, including BT_0154, BT_4000, BT_4562, BT_4563, BT_0328, and BT_1085, that reproducibly associate with components of the Lpt machinery, indicating that the *Bacteroides* Lpt system is more complex than the canonical machinery (Table 2). LptZ itself associates with multiple components of the Lpt complex and shows high-confidence interactions with the IM components LptF and LptG and the OM LptD-LptE complex (Fig.S5H). These interactions raise the possibility that LptZ simultaneously engages components at both membranes and adopts an extended conformation across the periplasm, potentially stabilized by one or more of the additional Lpt-associated proteins. Although this model remains speculative, it is structurally plausible, as we recently showed that other proteins in *B. thetaiotaomicron*, such as Dma1, can span the periplasm and connect the IM and OM^12^. The presence of these additional components also raises the possibility that the Lpt machinery does not exist as a single invariant complex but instead forms compositionally distinct assemblies with different substrate specificities. The canonical Lpt components may constitute a common transport core that associates with different combinations of LptZ and other accessory factors. In this model, LptZ-containing complexes could preferentially transport LES-containing SLPs, whereas LPS could be transported by complexes lacking LptZ, consistent with our finding that deletion of *lptZ* does not impair LPS transport. Alternatively, individual Lpt complexes may transport both LPS and SLPs, with substrate preference determined by their complement of associated factors. Defining the architecture and composition of these complexes will require dedicated structural and biochemical studies and represents an important direction for future work.

An important unresolved question is how SLP trafficking is mechanistically coupled to OMV biogenesis. Bacteroides OMVs are highly enriched in SLPs, and loss of LptZ simultaneously disrupts SLP trafficking and arrests OMV formation at the scission step. Because membrane curvature and budding still occur in ΔLptZ cells, SLPs do not appear to be required for the initiation of vesicle formation. Instead, their accumulation at the cell surface may facilitate a later step required for membrane constriction and scission. This could result from the collective effects of high SLP density on membrane organization, curvature, or protein crowding, or from the activity of one or more specific SLPs that promote vesicle release. Alternatively, the process of SLP translocation itself, rather than the presence of SLPs at the surface, may contribute to local remodeling of the OM during vesicle formation. Distinguishing among these possibilities will require determining whether restoration of individual SLPs, or of overall SLP abundance at the cell surface, is sufficient to restore OMV release in the absence of LptZ.

Overall, our findings identify LptZ as a *Bacteroidota*-specific component of the Lpt machinery and establish a mechanistic link between SLP trafficking and OMV biogenesis. We further suggest that specialization of a conserved envelope transport system has contributed to the remarkable diversification of the *Bacteroidota* cell surface. In gut-associated *Bacteroides*, SLPs are central to the degradation and acquisition of complex dietary and host-derived glycans, processes that strongly influence bacterial fitness and niche occupation within the intestinal microbiota^5^. By controlling the surface delivery and OMV packaging of these proteins, the LptZ-dependent pathway may therefore influence not only nutrient utilization by individual Bacteroides species but also their competitive fitness, interactions with other members of the microbiome, and ultimately the composition and function of the gut microbial community.

## MATERIALS AND METHODS

### Bacterial strains and growth conditions

Details of all bacterial strains, and plasmids used in this study are provided in Supplementary Table 3. *Escherichia coli* was cultured aerobically at 37 ⁰C in 2xYT or LB medium. *Bacteroides* strains were grown anaerobically at 37 ⁰C in a Coy anaerobic chamber (10% H₂, 5% CO₂, 85% N₂) in brain heart infusion (BHI) medium (Fisher Scientific) supplemented with 5 μg/ml hemin and 1 μg/ml vitamin K_3_. Where appropriate, antibiotics were added at the following concentrations: for *E. coli*, ampicillin (100 μg/ml), chloramphenicol (10 μg/ml), and kanamycin (50 μg/ml); for *Bacteroides thetaiotaomicron*, gentamicin (200 μg/ml), erythromycin (25 μg/ml), and tetracycline (10 μg/ml).

### Construction of deletion or insertion mutant in *Bt*

Gene deletion mutants were generated using the pSIE1 vector, as described previously ^36^. Briefly, ∼750 bp regions flanking the *bt_3341* (*lptZ*) were cloned into pSIE1. The resulting plasmids were introduced into *Bt* (WT) by conjugation. Transconjugants were selected on BHI agar supplemented with gentamicin and erythromycin. Counterselection was performed on BHI plates with or without 125 μg/ ml anhydrotetracycline (aTc). Candidate deletion mutants were screened by PCR and confirmed by whole-genome sequencing.

Plasmids were introduced into *Bt* by conjugation, as previously described. Transconjugants were selected on BHI agar supplemented with gentamicin to counter-select *E. coli* donors and erythromycin to identify positive transformants.

C-terminal 2xFLAG-tagged LptA and LptZ strains were generated using the pSIE1-mediated allelic exchange system described above. Briefly, approximately 1,000-bp regions upstream and downstream of the intended 2xFLAG insertion site, together with the sequence encoding the 2xFLAG epitope, were amplified and assembled into pSIE1. The resulting constructs were introduced into *B. thetaiotaomicron*, and the 2xFLAG sequence was inserted in-frame at the C terminus of LptA or LptZ following the allelic exchange procedure described above.

### Isolation and Purification of TM, IM, OM, and OMVs

Total membranes (TM), inner membranes (IM), outer membranes (OM), and outer membrane vesicles (OMVs) were purified as described previously ^7,9^, with minor modifications. *Bt* strains from BHI agar plates were initially inoculated into 3 mL of BHI broth containing appropriate antibiotics (if required) and incubated anaerobically at 37 ⁰C overnight. The following day, cultures were diluted 1:50 into 50 mL fresh BHI medium and incubated anaerobically at 37 ⁰C for ∼16 h. Cells were harvested by centrifugation at 6000 x g for 10 min at 4 ⁰C. The supernatant was used for OMV isolation, while the cell pellet was reserved for membrane fractionation. Supernatants were passed through 0.22 μm filter units (Stericup Quick Release, Millipore) to remove residual cells and debris. The clarified culture supernatant (∼50 mL) was subjected to ultracentrifugation at 200,000 x g for 2 h at 4 ⁰C using a Ti-45 rotor (Optima L-100 XP, Beckman Coulter). Resulting OMV pellets were resuspended in 200 μL of 50 mM HEPES buffer (pH 7.4) per 50 OD_600_ units of culture.

For membrane isolation, the cell pellet was resuspended in 20 mL of 50 mM HEPES buffer (pH 7.4) supplemented with EDTA-free protease inhibitors (Thermo Scientific). Cells were lysed by two passes through a Constant Systems cell disruptor at 35 kpsi. The lysate was centrifuged at 8,000 x g for 10 min at 4 ⁰C to remove unbroken cells and debris. Total membranes were collected by ultracentrifugation at 200,000 x g for 1 h at 4 ⁰C. The resulting TM pellet was resuspended in 1 mL of 50 mM HEPES (pH 7.4) per 50 OD_600_ units of starting culture.

To separate IM and OM, TM suspensions were incubated with 50 mM HEPES (pH 7.4) containing 1% (w/v) N-lauroyl sarcosine in 1.5 mL tubes with gentle rocking at room temperature for 1 h. Samples were centrifuged at 100,000 x g for 1 h at 4⁰C. The supernatant, containing solubilized IM, was collected, while the pellet, containing OM, was resuspended in 50 mM HEPES (pH 7.4). Isolated fractions were then lyophilized for downstream proteomic analysis.

### Localization of LptZ as detected by WB

To assess the subcellular localization of LptZ, the *lptZ* gene and BT_4362 (SecA homolog, an inner membrane protein used as a control) were cloned into plasmid pWW3452 ^34^, each with a C-terminal 3xFLAG epitope tag. The resulting constructs were introduced into *Bt* strains: pWW3452:LptZ into a Δ*lptZ* background, and pWW3452:BT_4362 into WT *Bt* by conjugation as described previously. TM, IM, and OM were isolated from each strain as described above. Equivalent amounts of protein from each membrane fraction were mixed with SDS sample buffer and resolved on 12% SDS-PAGE gels. FLAG-tagged LptZ and BT_4362 were detected using mouse anti-FLAG primary antibody (1:1000; Sigma) and IRDye-conjugated donkey anti-mouse secondary antibody (1:10000; LI-COR), followed by imaging on an Odyssey CLx system (LI-COR Biosciences). For verification of membrane fraction purity, the OM marker protein SusC was detected in the same blots using rabbit anti-SusC (1:1000) primary antibody and IRDye-conjugated goat anti-rabbit secondary antibody (1:10000; LI-COR).

### OM-stress response assay

To assess OM integrity, *Bt* strains were grown overnight at 37 ⁰C under anaerobic conditions in BHI broth. Cultures were adjusted to an OD_600_ of 1.0 in fresh BHI and subsequently subjected to 10-fold serial dilutions in sterile 96-well microtiter plates. From each dilution, 10 µL was spotted onto BHI agar plates supplemented with either 1 mM EDTA, 50 µg/mL vancomycin, or 10 µg/mL polymyxin B. Plates lacking any stressors served as growth controls. Spotted plates were incubated anaerobically at 37 ⁰C for 48 h. Assays were performed in three independent biological replicates. One representative experiment was selected for the figure presentation.

### Growth assay with *Bt* strains

*Bt* WT and Δ*lptZ* strains were cultured overnight at 37 ⁰C under anaerobic conditions in BHI broth. Overnight cultures were centrifuged at 6000 x g for 10 min and washed twice with minimal medium (MM) lacking carbohydrate supplementation. The optical density at 600 nm (OD_600_) of the resulting suspensions was measured and normalized to OD_600_ =0.1 in MM supplemented with the indicated carbon sources (0.5% w/v final concentration). Growth assays were conducted in sterile, round-bottom 96-well polystyrene microplates. Cultures were incubated anaerobically at 37 ⁰C under static conditions. OD_600_ readings were recorded every 30 min using a Smart Reader 96-T (Accuris Instruments) following 10 seconds of orbital shaking to ensure homogenization. Each growth condition was tested in technical triplicate and independently repeated in three biological replicates.

### Sus protein induction in the presence of starch and detection by western blot

*Bt* WT, Δ*lptZ*, and complemented strains were cultured overnight at 37 ⁰C under anaerobic conditions. The following day, cultures were subcultured 1:50 into fresh BHI broth, with or without 0.5 % (w/v) starch, and incubated anaerobically for 24 h. Following incubation, OD_600_=1.0 cells were harvested by centrifugation, and resuspended in protein dye buffer. Samples were boiled for cell lysis and total protein extraction. Proteins were resolved by SDS-PAGE on 12% acrylamide gels and transferred to nitrocellulose membranes. Membranes were probed with rabbit polyclonal antibodies against SusD, SusE, SusF, and SusG (1:1000 dilution). IRDye-conjugated goat anti-rabbit secondary antibodies (1:10,000; LI-COR) were used for detection.

### Affinity purification of LptZ and identification of interacting partners in *Bt*

To identify interacting partners of LptZ, a dual-tagged construct encoding *lptZ* with C-terminal His_10_ and 3xFLAG tags was cloned into the pWW3515 expression vector ^34^. The resulting plasmid (pWW3515:LptZ) was introduced into a *Bt* Δ*lptZ* strain, where the tagged LptZ was verified to be functionally active. WT strain served as a negative control. WT and Δ*lptZ*:: pWW3515:LptZ-His_10_-FLAG(3X) were grown overnight in BHI medium. Cultures were diluted 1:50 into 800 mL fresh BHI and incubated anaerobically for an additional 16–18 h. Cells were harvested by centrifugation at 10000g for 20 min at 4 ⁰C, washed with 0.9% NaCl, and resuspended in TBS buffer (40 mM Tris-HCl, pH 8.0, 150 mM NaCl) supplemented with EDTA-free protease inhibitor cocktail (Thermo Scientific). Cells were lysed using a Constant Systems cell disruptor at 30,000 psi (two passes). To degrade genomic DNA, 25 μL DNase I was added, and the lysate was incubated at 4 ⁰C for 30 min. Total membrane fractions were isolated by ultracentrifugation at 200,000 x g for 1 h at 4 ⁰C. Membrane pellets were resuspended in 20 mL TBS (40mM Tris pH8.0, 150mM NaCl) containing 1% n-dodecyl-β-D-maltoside (DDM; Thermo Scientific) and incubated overnight at 4 ⁰C with gentle agitation to solubilize membrane proteins. Insoluble debris was removed by a second ultracentrifugation (200,000 × g, 1 h, 4 ⁰C), and the supernatant containing solubilized proteins was collected. Affinity purification was performed in two steps. First, the solubilized protein solution was incubated with 500 μL Ni-NTA resin (equivalent to 1 mL of 50% slurry) for 1 h at 4 ⁰C with constant rotation. Resin was transferred to a gravity-flow column (Bio-Rad), washed 10 times with 2 mL of wash buffer (TBS + 1% DDM + 40 mM imidazole), and eluted with 4 sequential 1 mL fractions of elution buffer (TBS + 1% DDM + 500 mM imidazole). To remove excess imidazole, eluted fractions were pooled, concentrated to 500 μL using a 30 kDa cutoff Vivaspin concentrator (Sartorius), and this step was repeated twice with 10-fold dilution in ice-cold TBS containing 1% DDM each time, and finally reconcentrated to ∼500 μL. For FLAG-based purification, the concentrated eluate was incubated with 100 μL anti-DYKDDDDK (FLAG) affinity resin (Thermo) for 1 h at 4 ⁰C in 1.5 mL microcentrifuge tubes. The resin was washed four times with 500 μL TBS, and bound protein complexes were eluted in four sequential 200 μL fractions using 0.1 M glycine (pH 2.8). Eluates were pooled, lyophilized, and submitted for proteomic analysis. All purification steps were performed at 4 ⁰C using buffers supplemented with EDTA-free protease inhibitors (Thermo Scientific).

To identify protein interactions that may be weak or transient, *in vivo* protein complexes containing LptZ or LptA were stabilized using the membrane-permeable crosslinker dithiobis(succinimidyl propionate) (DSP). *Bt* strains carrying genomically encoded C-terminal 2xFLAG-tagged LptZ or LptA were generated as described previously. The WT *Bt* strain was used as a negative control. Strains were grown as described above, and 500 mL of overnight culture was harvested by centrifugation and washed twice with PBS to remove residual medium. Cell pellets were resuspended in 50 mL PBS, and DSP was added to a final concentration of 0.5 mM. Samples were incubated for 30 min to allow crosslinking of protein complexes. The reaction was quenched by addition of Tris-HCl (pH 7.5) to a final concentration of 100 mM. Cells were subsequently harvested by centrifugation, washed with TBS2 buffer (50 mM Tris-HCl, pH 8.0, 150 mM NaCl), and resuspended in 50 mL of the same buffer.

Cells were lysed, and total membrane fractions were isolated and solubilized as described above. LptZ-2xFLAG or LptA-2xFLAG complexes were purified from the solubilized membrane fraction using 200 μL anti-DYKDDDDK (FLAG) affinity resin as described above. The purified complexes were subsequently analyzed by proteomics.

### Analysis of LptA- and LptZ-dependent association with LES-containing lipoproteins

To determine whether LptA and LptZ preferentially associate with lipoproteins containing an N-terminal lipoprotein export specificity (LES) sequence, three non-LES lipoproteins (BT_0705, BT_2548, and BT_4399) and a chimeric form of BT_2548 containing the LES sequence form SusD [BT_2548(LES)] were cloned into pWW3452 ^34^ as C-terminal His10-tagged proteins. The resulting constructs were introduced into WT *Bt* and strains carrying genomically encoded C-terminal 2xFLAG-tagged LptA or LptZ, respectively.

Strains were grown as described above, and 400 mL cultures were treated with DSP at a final concentration of 1 mM for 1 h to stabilize protein-protein interactions. LptA- or LptZ-containing complexes were purified by FLAG affinity chromatography as described above, except that bound proteins were eluted directly by addition of 1x SDS-PAGE sample buffer to the resin. Eluted proteins were separated by SDS-PAGE and analyzed by immunoblotting. LptA and LptZ were detected using an anti-FLAG antibody, whereas the His10-tagged non-LES lipoproteins (BT_0705, BT_2548, BT_4399, and BT_2548(LES)) were detected using an anti-His antibody. SusD and SusF were detected using protein-specific antibodies as described in previous sections.

#### Sample preparation for Proteomic analysis

Lyophilized membrane fractions and immunoprecipitated protein samples were solubilized in 4% SDS, 100 mM HEPES by boiling for 10 min at 95 ⁰C. For membrane fractions, protein concentrations were assessed by bicinchoninic acid protein assay (Thermo Fisher Scientific) and 100 µg of each biological replicate was prepared for proteomic analysis using S-Trap mini columns (Protifi, USA) according to the manufacturer’s instructions. Immunoprecipitated protein samples were prepared using S-Trap mini columns without protein quantification, assuming ∼20 µg total proteins. All samples were reduced with 10 mM Dithiothreitol for 10 min at 95 ⁰C and then alkylated with 40 mM Iodoacetamide in the dark for 1 h. Samples were acidified to 1.2% phosphoric acid and diluted with seven volumes of S-Trap wash buffer (90% methanol, 100 mM Tetraethylammonium bromide pH 7.1) before being loaded onto S-Traps and washed 3 times with 400 µL of S-Trap wash buffer. Samples were then digested with 2 µg of Trypsin/Lys-C (Promega, 1:50-1:10 protease/protein ratio) in 100 mM Tetraethylammonium bromide overnight at 37 °C before being collected by centrifugation with washes of 100 mM Tetraethylammonium bromide, followed by 0.2% formic acid, followed by 0.2% formic acid / 50% acetonitrile. Samples were dried down and further cleaned up using C18 Stagetips ^1,2^ to ensure the removal of any particulate matter.

#### Reverse phase Liquid chromatography–mass spectrometry

Prepared C18 purified peptides from each sample were re-suspended in Buffer A* (2% acetonitrile, 0.01% trifluoroacetic acid) and separated using a two-column chromatography setup composed of a PepMap100 C_18_ 20-mm by 75-μm trap and a PepMap C_18_ 500-mm by 75-μm analytical column using a Dionex Ultimate 3000 UPLC (Thermo Fisher Scientific). Each sample was concentrated onto the trap column at 5 μl/min for 6 min with Buffer A (0.1% formic acid, 2% DMSO) before being subjected to analytical separation at 300 nl/minute. Immunoprecipitated samples were analysed on either an Orbitrap Exploris 480 (non-cross-linked samples) or an Orbitrap Eclipse (cross-linked samples) using 59-minute analytical gradients by altering the buffer composition from 2% Buffer B (0.1% formic acid, 77.9% acetonitrile, 2% DMSO) to 22% B over 29 min, then from 22% B to 40% B over 10 min, then from 40% B to 80% B over 5 min. The composition was held at 80% B for 5 min, and then dropped to 2% B over 2 min before being held at 2% B for another 8 min. The Orbitrap Exploris 480 and Eclipse Mass Spectrometers were operated in a data-dependent switching between a single Orbitrap MS scan (300-1600 m/z and a resolution of 120k) was acquired every 3 seconds followed by Orbitrap MS/MS HCD scans of precursors (fixed NCE 30%, maximal injection time of 50 ms and a resolution of 30k). Membrane fractions samples were analysed using 89-minute analytical gradient on an Orbitrap Ascend by altering the buffer composition from 2% Buffer B (0.1% formic acid, 77.9% acetonitrile, 2% DMSO) to 22% B over 59 min, then from 22% B to 40% B over 10 min, then from 40% B to 80% B over 5 min. The composition was held at 80% B for 5 min, and then dropped to 2% B over 2 min before being held at 2% B for another 8 min. The Orbitrap Ascend Mass Spectrometer was operated in a data-dependent switching between a single Orbitrap MS scan (300-1350 m/z and a resolution of 120k) was acquired every 3 seconds followed by Orbitrap MS/MS HCD scans of precursors (fixed NCE 25%, maximal injection time of 27 ms and a resolution of 15k).

#### Proteomic data analysis

Identification and LFQ analysis were accomplished using MaxQuant (v2.4.7.0)^3^ using the *Bacteroides thetaiotaomicron* VPI-5482 proteome (Uniprot: UP000001414), allowing for oxidation as a variable modification of Methionine and carbamidomethylation of cysteine set as a fixed modification. A Trypsin specificity was set with the LFQ and “Match Between Run” enabled to allow comparison between samples. The resulting data files were processed using Perseus (v1.4.0.6)^4^ with missing values imputed based on the observed total peptide intensities with a range of 0.3σ and a downshift of 1.8σ prior to comparison of groups using student t-tests.

To identify OMV-specific proteins, we compared the proteomes of the OM and OMVs isolated from WT *Bt*. Proteins with a log_2_ fold change (OM/OMV) ≤ -1.5 were classified as OMV-specific, as these were considered highly enriched in OMVs relative to the OM (Dataset 1). Ribosomal proteins were annotated based on their predicted assignment to the 30S or 50S ribosomal subunits, while other cytosolic proteins were designated according to their predicted cellular compartment (Dataset 1).

For comparison of IM and OM proteomes, we determined the relative IM and OM abundance of proteins in WT and ΔLptZ strains, expressed as the log₂(IM/OM) ratio (Dataset 3). To validate the efficiency of membrane fractionation, proteins were first examined according to their predicted cellular localization, ensuring that IM and OM proteins were appropriately enriched in their respective fractions.

To classify lipoproteins into LES and non-LES groups, we first selected lipoproteins that were not transcriptionally altered (-1 < Log_2_ fold change < 1). Among these, lipoproteins enriched in OMVs relative to the OM, defined by Log_2_(OM/OMV) < -1.5, were considered OMV-specific, whereas those with Log₂(OM/OMV) >1 were classified as OM-specific. Lipoproteins with intermediate enrichment (-1.5 < Log_2_(OM/OMV) < 1) were not considered highly enriched in either fraction. For OMV- and OM-enriched lipoproteins, sequence logos were generated using the ten residues immediately following the conserved cysteine. Lipoproteins with intermediate enrichment (-1.5 < Log_2_(OM/OMV) < 1) were further divided into LES and non-LES groups based on the presence of the corresponding sequence motif (Datasets 9-11).

To assess whether the absence of LptZ or LolB affects lipoprotein transport, we compared the IM/OM distribution between WT and ΔLptZ or LolB strains by calculating log_2_[(IM/OM of ΔLptZ)/(IM/OM of WT)] or log_2_[(IM/OM of LolB-)/(IM/OM of WT)], respectively. Positive values indicate a higher IM/OM ratio in ΔLptZ or LolB- relative to WT, consistent with impaired transport of the respective protein to the OM. Positive values denote a relative increase in IM/OM ratio in ΔLptZ compared to WT, suggesting impaired localization of the protein to the OM.

To assess the lipoproteins affected by LolB depletion, we obtained the published proteomics dataset from Armbruster et al^33^. The dataset was reanalyzed independently for the purposes of this study to identify lipoproteins whose abundance was affected between WT and LolB-depleted *Bt*. Protein abundance data and associated statistical values were extracted from the published dataset and analyzed.

For multi-comparison analyses (e.g., Fig. 5A-E; Fig.S12A-C; Datasets 8-13), we included all proteins for which at least one comparison: Log_2_(IM/OM) in WT, Log_2_(IM/OM) in ΔLptZ, Log_2_(IM/OM) in LolB- or Log_2_(OM) between WT and ΔLptZ-was statistically significant.

### Co-expression and pull-down of LptZ interacting proteins

To assess interactions between LptZ and candidate partner proteins, the *lptZ* coding sequence was cloned into the pBAD33.1 ^37^ (Addgene) vector with a C-terminal FLAG tag. Putative interacting partners, including LptD (BT_2133), LptE (BT_4376), LptA (BT_3849), LptC (BT_4369), LptF (BT_2417), LptG (BT_0834), and LptB (BT_3837), were cloned into the pBAD-Myc-His vector (invitrogen) with C-terminal His_10_ tags. Each pBAD-Myc-His construct was co-transformed with either pBAD33.1:LptZ-FLAG or empty pBAD33.1 into *E. coli* BL21(DE3). Overnight cultures of transformed strains were diluted 1:100 into 50 mL 2xYT medium containing ampicillin and chloramphenicol, and grown at 37 ⁰C with shaking (200 rpm). After 2 h, protein expression was induced with 0.1% (w/v) L-arabinose, followed by an additional 4 h incubation at 37 ⁰C with agitation (160 rpm).

Cells were harvested and resuspended in 5 mL TBS buffer (40 mM Tris-HCl, pH 8.0, 150 mM NaCl) containing 1% n-dodecyl-β-D-maltoside (DDM; Thermo Scientific). Lysis was performed using a Constant Systems cell disruptor at 30,000 psi (two passes), followed by 1 h incubation at 4 ⁰C with rotation to solubilize membrane proteins. For input analysis, 100 μL of lysate was mixed with 100 μL 2 x SDS-PAGE loading buffer, boiled at 95 ⁰C for 10 min, and stored. For pull-down assays, 500 μL of the lysate was incubated with 50 μL anti-DYKDDDDK (FLAG) affinity resin (Thermo) for 1 h at 4 ⁰C with rotation. The mixture was transferred to a gravity-flow column and washed ten times with 500 μL TBS containing 1% DDM. Bound protein complexes were eluted with 200 μL 1x SDS-PAGE sample buffer, incubated at 95 ⁰C for 5 min, and centrifuged to pellet the resin. The supernatant was collected as the output fraction.

Both input and output samples were resolved by SDS-PAGE on 12% acrylamide gels and transferred to nitrocellulose membranes. FLAG-tagged LptZ was detected using mouse anti-FLAG monoclonal antibodies (Sigma), while His-tagged candidate partners were probed using rabbit anti-His polyclonal antibodies (Sigma). Appropriate secondary antibodies and infrared imaging were used as described above. All experiments were independently repeated at least three times, with representative data shown.

### Bacterial two-hybrid assay

Protein-protein interactions were evaluated using the bacterial adenylate cyclase two-hybrid (BACTH) system ^38^. The *lptZ* gene (BT_3341) was cloned into the pUT18 vector, while candidate interacting genes were cloned into the compatible pKNT25 vector. All constructs were designed with C-terminal fusions of the T18 or T25 fragments of *Bordetella pertussis* adenylate cyclase to preserve the native N-terminal signal sequences of LptZ and its putative interacting partners, as these proteins are membrane-associated.

Recombinant plasmid pairs were co-transformed into *E. coli* BTH101 reporter cells. Three independent colonies from each transformation were cultured overnight in 2xYT medium supplemented with ampicillin, kanamycin, and 0.5 mM IPTG at 30 ⁰C. The next day, 5 μL of each culture was spotted onto LB agar plates containing ampicillin, kanamycin, 40 μg/mL X-gal, and 0.5 mM IPTG. Plates were incubated at 30 ⁰C for 24 h, and protein interactions were inferred from the development of blue coloration. Each experiment was performed in triplicate with consistent results; representative data are shown.

### LPS Isolation and Quantification

LPS was isolated from various cellular and vesicular fractions, including TM, IM,OM, and OMVs. Each fraction was treated with proteinase K and incubated at 37 ⁰C for 4 h to digest contaminating proteins. For TM and OMV fractions, the loading amounts were normalized based on the OD_600_ of the originating cultures. For comparative analyses of IM, OM, and TM fractions, samples were normalized by total protein concentration as determined by a standard protein assay. LPS samples were separated on 10-20% SDS-PAGE gels (Invitrogen) and quantified using two complementary methods:

Silver Staining: Gels were stained using a modified protocol of Tsai and Frasch ^39^. Briefly, after electrophoresis, gels were fixed overnight in 200 mL of 40% ethanol and 5% acetic acid. Oxidation was carried out for 5 min in 100 mL of 0.7% periodic acid prepared in the same fixative. Gels were washed three times for 15 min each with Milli-Q water, then incubated in the dark for 10 min in a staining solution composed of 28 mL 0.1 M NaOH, 2 mL NH₄OH, 5 mL 20% AgNO₃, and 115 mL Milli-Q water. After three additional washes, gels were developed in 200 mL Milli-Q water containing 10 mg citric acid and 100 μL formaldehyde.

Western blot: Alternatively, LPS was detected using a custom-made rabbit anti-*Bt* LPS antibody. Following SDS-PAGE, LPS was transferred to a nitrocellulose membrane, probed with the primary anti-LPS antibody, and detected using a corresponding IRDye-conjugated secondary antibody. Signal was visualized using a LI-COR Odyssey infrared imaging system, as previously described.

### Cryo-ET sample preparation and imaging

*Bt* cells were grown on BHI-agar plates as described previously. Subsequently, cells were collected from the plate using a loop and resuspended in 1 mL of 1x PBS to a final OD_600_ of 3.0 for cryo-ET experiments.

R2/2 carbon-coated 200 mesh copper Quantifoil grids (Quantifoil Micro Tools) were glow-discharged for 30 seconds. Then, cells were mixed with a solution of 10-nm gold beads treated with Bovine Serum Albumin. 4 μl of this mixture was applied on the grids in a Vitrobot chamber (FEI). Subsequently, the extra fluid was blotted off using a Whatman filter paper in the Vitrobot chamber with 100% humidity and the grids were plunge-frozen in a cryogen (liquid ethane).

Samples imaging was performed at the Advanced Electron Microscopy at the University of Chicago. Cells were imaged using a Titan Krios transmission electron microscope operating at 300 kV and equipped with a BioQuantum K3 imaging filter (Gatan). Data were collected using Tomography 5 software with each tilt series ranging from -60° to 60° in 3° increments and an underfocus of ∼6-8 μm and a total dose of 130 e^-^/Å^2^.

### Image processing and data visualization

Three-dimensional reconstructions of tilt-series were performed using the IMOD software package^40^. Tomograms segmentations were done using Dragonfly software (https://www.theobjects.com/dragonfly/index.html). A series of TIFF files were generated from an MRC tomogram file using “mrc2tif” command in IMOD software package. The generated TIFF image stack was imported to Dragonfly followed by intensity scale calibration. Initial training model was created in the Segmentation Wizard for deep learning. At least three representative tomogram slices were selected to train the model. Regions of interest, such as outer and inner membranes, were labeled as different color classes, and the rest of unlabeled region was recognized as background. The AI-driven algorithm constructs the 3D model through tracing the 2D gray contrast of each tomogram slice. The generated model was then modified manually to remove the noise and fill in the missing parts. Smooth Mesh tool was applied for polishing the model surface.

### Phylogenetic analysis of LptZ

The LptZ protein sequence (BT_3341) was queried against the RefSeq Protein database (version 2024-10-13) using the BLASTp tool via the Galaxy Europe platform (https://usegalaxy.eu/). Search parameters were set to an E-value threshold of 0.001 and a minimum query coverage of 80%. This yielded 5,107 putative LptZ homologs across 2,245 unique species. From this set, the SILVA database was searched for the 16S rRNA gene sequences of matching bacterial species and successfully extracted 1028 sequences ^41^. 213 additional 16S rRNA sequences were retrieved from NCBI (ncbi.nlm.nih.gov), using annotation files and the assembled genomes of the corresponding NCBI entries, leading to a total of 1,241 sequences. The 16S rRNA sequences were aligned using MEGA11^42^, and a phylogenetic tree was subsequently constructed and visualized using iTOL (Interactive Tree of Life)^43^.

### RNA Sequencing Sample Collection, Library Preparation, and Analysis

WT and Δ*lptZ* strains were grown overnight in BHI medium and then diluted to an OD_600_ of 0.1 in 10 mL fresh medium. Cultures were incubated anaerobically at 37 ⁰C for 4 h. Four biological replicates were prepared for each strain. Following incubation, cultures were normalized, and a volume equivalent to an OD_600_ of 4.0 was pelleted by centrifugation at 8,000 x g for 90 seconds. Cell pellets were resuspended on ice in 1 mL of TRIzol reagent (Invitrogen) containing 10 μL of 5 mg/mL glycogen. Samples were flash-frozen and stored at −80 ⁰C until RNA extraction. For extraction, samples were thawed on ice, centrifuged, and the supernatant was treated with chloroform. RNA was isolated from the aqueous phase using the RNeasy Mini Kit (Qiagen). RNA quality was assessed via agarose gel electrophoresis and A_260_/A_280_ measurements. Extracted RNA was stored at −80 ⁰C with SUPERase-In RNase inhibitor (Life Technologies) until library preparation.

RNA sequencing (RNA-seq) was performed as previously described ^44^. Briefly, 400 ng of total RNA per sample was used to generate cDNA libraries following the RNAtag-Seq protocol. Libraries were PCR-amplified and sequenced using an Illumina NextSeq500 platform, yielding over 7 million reads per sample. Data analysis was performed using our in-house pipeline, Aerobio ^45^. Raw reads were demultiplexed based on 5′ and 3′ indices, trimmed to 59 base pairs, and quality filtered (96% of bases with Q > 14). Filtered reads were mapped to the corresponding reference genomes using Bowtie2 with the -very-sensitive option (-D 20 –R 3 –N 0 –L 20 –i S,1,0.50). Mapped reads were quantified using featureCounts, and differential expression analysis was performed with DESeq2 ^12,44^. Genes were considered differentially expressed if they met both criteria: |log₂ fold change| > 1 and adjusted p-value (padj) < 0.05. All comparisons were made between WT and Δ*lptZ* samples under the conditions described above. Reproducibility across biological replicates was confirmed by high Spearman correlation (R > 0.95).

## Data availability

The mass spectrometry proteomics data has been deposited in the Proteome Xchange Consortium via the PRIDE partner repository with the data set identifier: PXD065577 [Username: Password: 0mHFJ9zPNWbA], PXD065584 [Username: Password: FZWsAcYwQvuK] and PXD083240 [Project accession: PXD083240 and Token: TEPZFK21RHJM]. Raw RNA-Seq datasets are available at the Sequence Read Archive (BioProject accession number PRJNA1284933) (https://submit.ncbi.nlm.nih.gov/subs/sra/SUB15428424/overview).

## Author contributions

B.J., M.F.F., M.K., and E.J.P. designed research; B.J., E.J.P., T.Z., M.G.S., M.J.M., N.E.S., J.C.O.-M., M.N., and M.K., performed research; B.J., T.Z., M.K., M.J.M., N.E.S., J.C.O.-M., and M.F.F. analyzed data; and B.J. and M.F.F. wrote the paper. All authors edited and approved the manuscript.

## Acknowledgements

We thank all members of the Feldman lab for critically reading the manuscript. This work was supported by NIH grants R01AI181213 and R21AI168719 to M.F.F. and R01AI148470 to TVO and JOM., N.E.S is supported by an Australian Research Council Future Fellowship (FT200100270) and an ARC Discovery Project Grant (DP210100362) and a National Health and Medical Research Council Ideas grant (2018980). We thank the Melbourne Mass Spectrometry and Proteomics Facility of The Bio21 Molecular Science and Biotechnology Institute for access to MS instrumentation. Research in the Cava laboratory is supported by the Swedish Research Council, the Laboratory for Molecular Infection Medicine Sweden (MIMS), Umeå University, the Knut and Alice Wallenberg Foundation (KAW), and the Kempe Foundation. We thank Lance Bottini for his diligent care of our instruments.

## Competing interests

The authors declare no competing interests.

## Declaration of generative AI and AI-assisted technologies in the manuscript preparation process

During the preparation of this work, the author(s) used ChatGPT for English-language editing. The author(s) reviewed and edited the output as needed and take full responsibility for the content of the published article.

---

**Figure S1:**
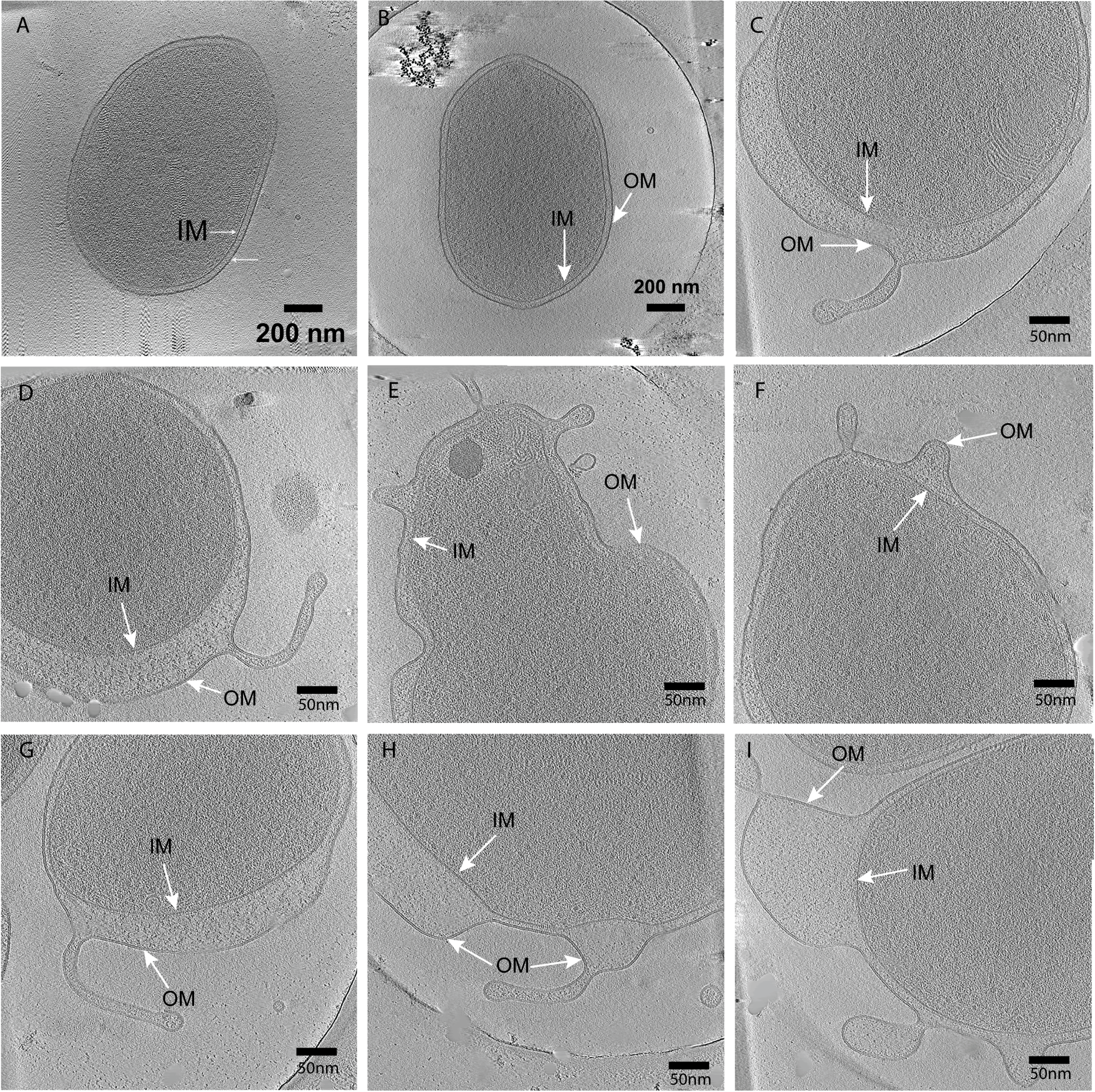
Slice through cryo-electron tomograms of WT (A-B) and Δ*lptZ* mutant cells (C-I) highlighting the irregular morphologies of mutant cells including extended periplasmic space and the presence of tubular structures.

**Figure S2:**
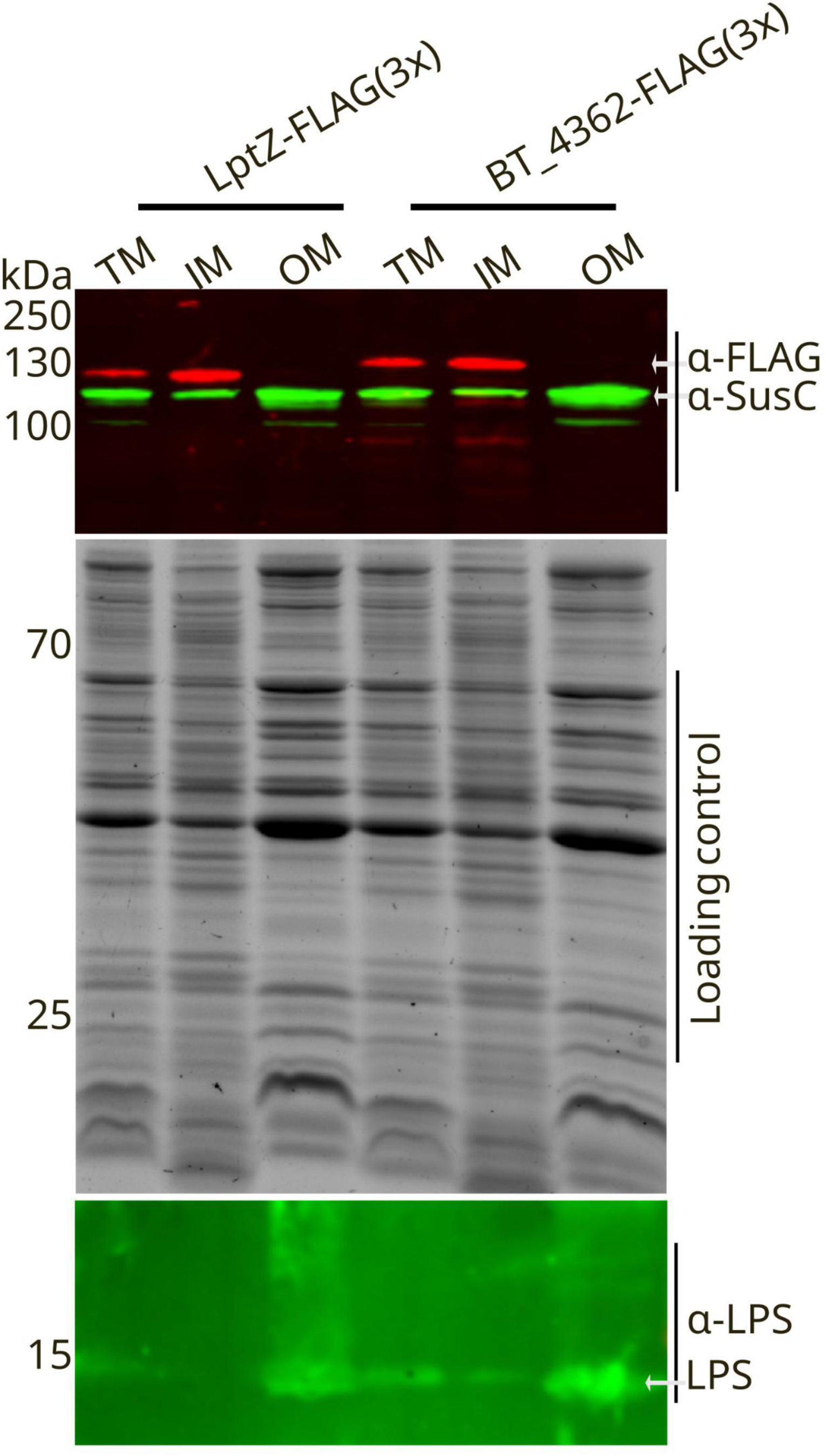
LptZ localizes to the IM. **Upper panel**. Western blot of FLAG-tagged LptZ and SecA (BT_4362) expressed in WT. LptZ shows IM localization predominantly. SusC (BT_3702) is found to be enriched in the OM fraction and serves as an OM marker. **Middle panel.** SDS-PAGE confirms equal loading. **Lower panel**. Western blot of proteinase K-treated TM, IM, and OM samples, probed with an anti-LPS antibody, reveals a specific LPS band (arrow) used as an OM control.

**Figure S3:**
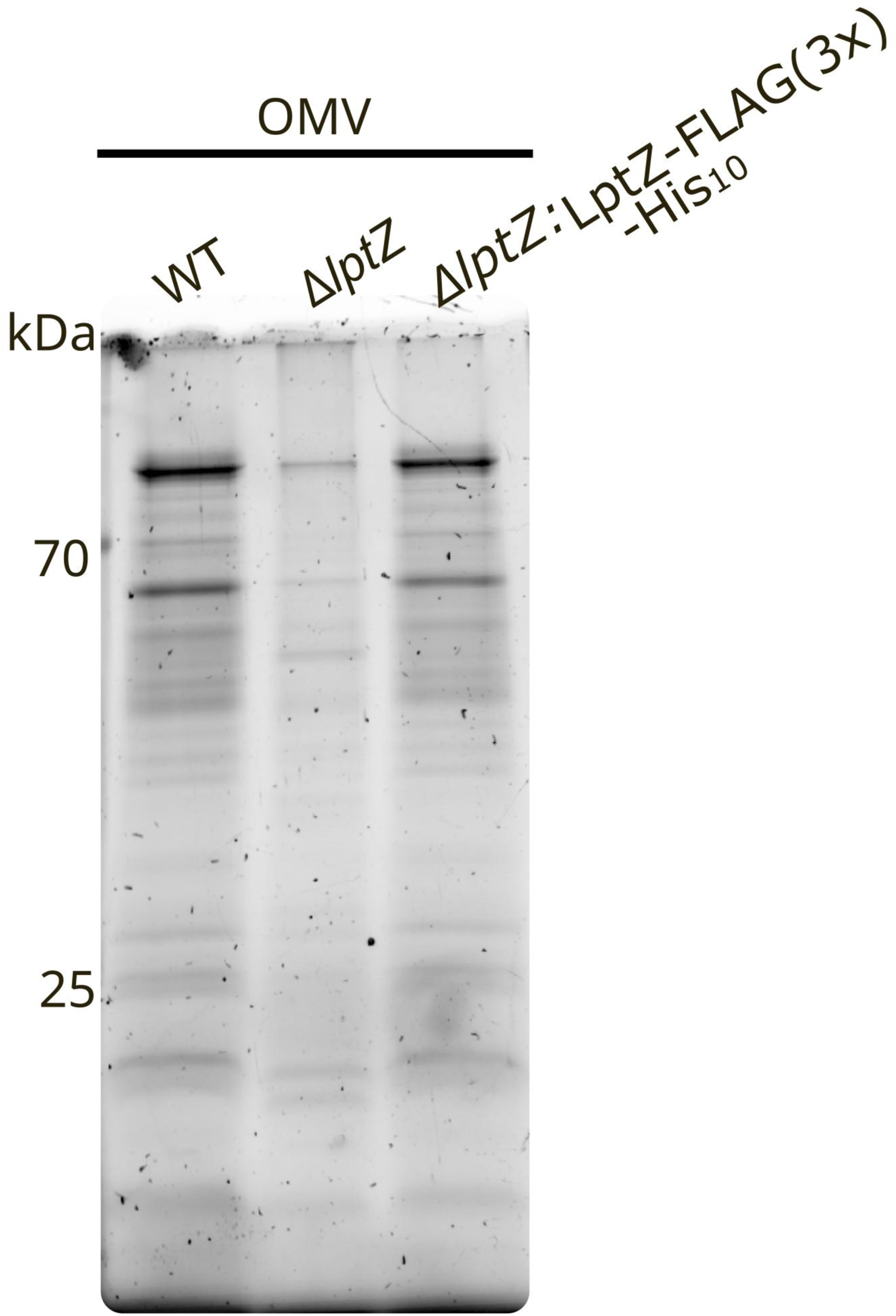
Dual-tagged (3xFLAG-His₁₀-tagged) LptZ retains its activity. OMVs were purified from WT, Δ*lptZ* mutant, and complemented (LptZ-FLAG(3x)-His10) strains and subjected to SDS-PAGE analysis. The similar OMV protein profiles of the WT and complemented strains indicate that the dual-tagging of LptZ does not affect its activity.

**Figure S4.**
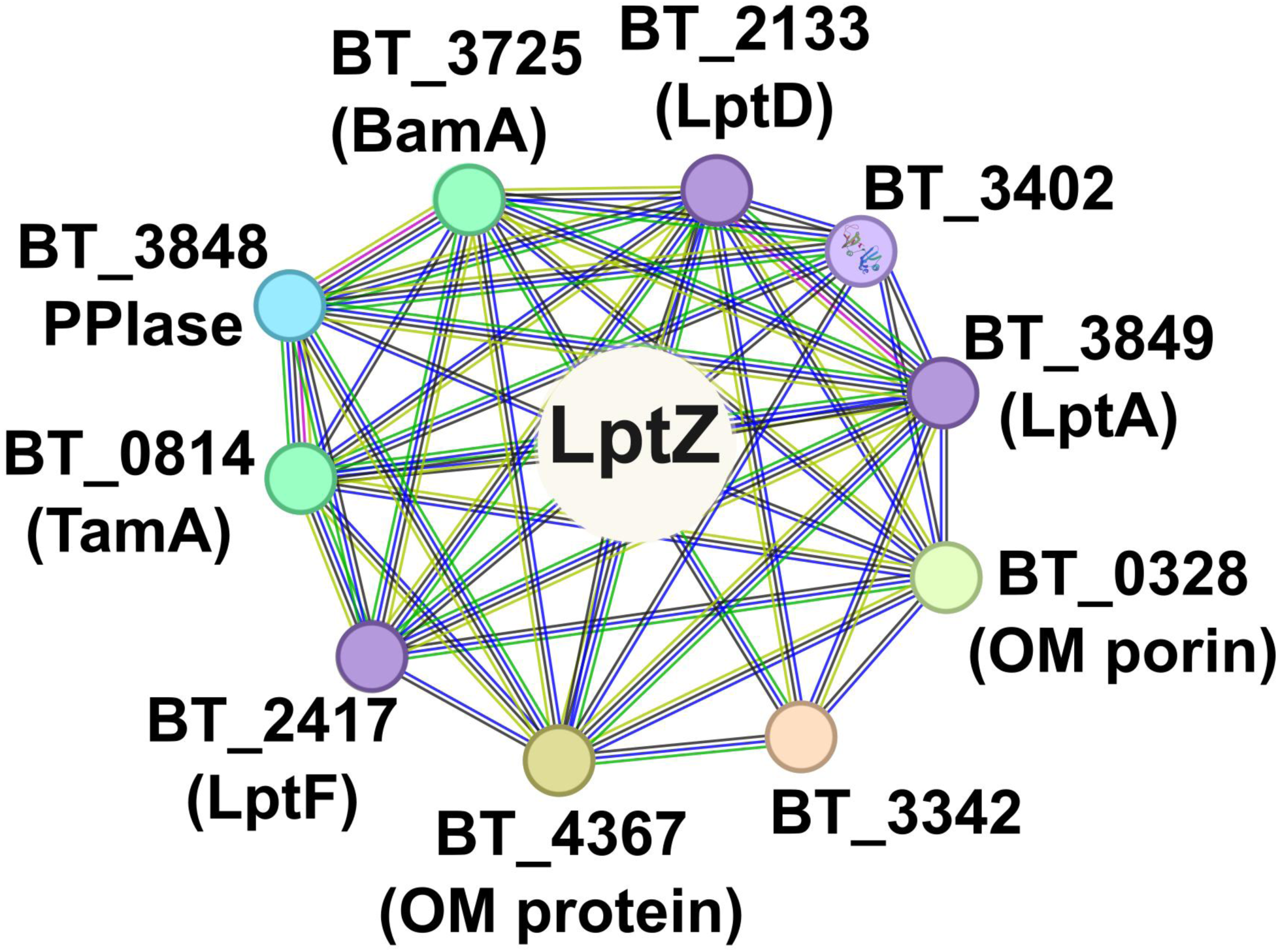
STRING-based interaction network of LptZ reveals predicted associations with LPS transport components (LptA [BT_3849], LptD [BT_2133], LptF [BT_2417]), β-barrel assembly factors (BamA [BT_3725], TamA [BT_0814]), and other OM-associated proteins (BT_3848, BT_0328, BT_4367). Lines indicate associations between proteins: Blue: known interactions from curated databases; Pink: experimentally determined; Green: gene neighborhood; Red: gene fusions; Light blue: text mining; Black: co-expression; Purple: protein homology.

**Figure S5:**
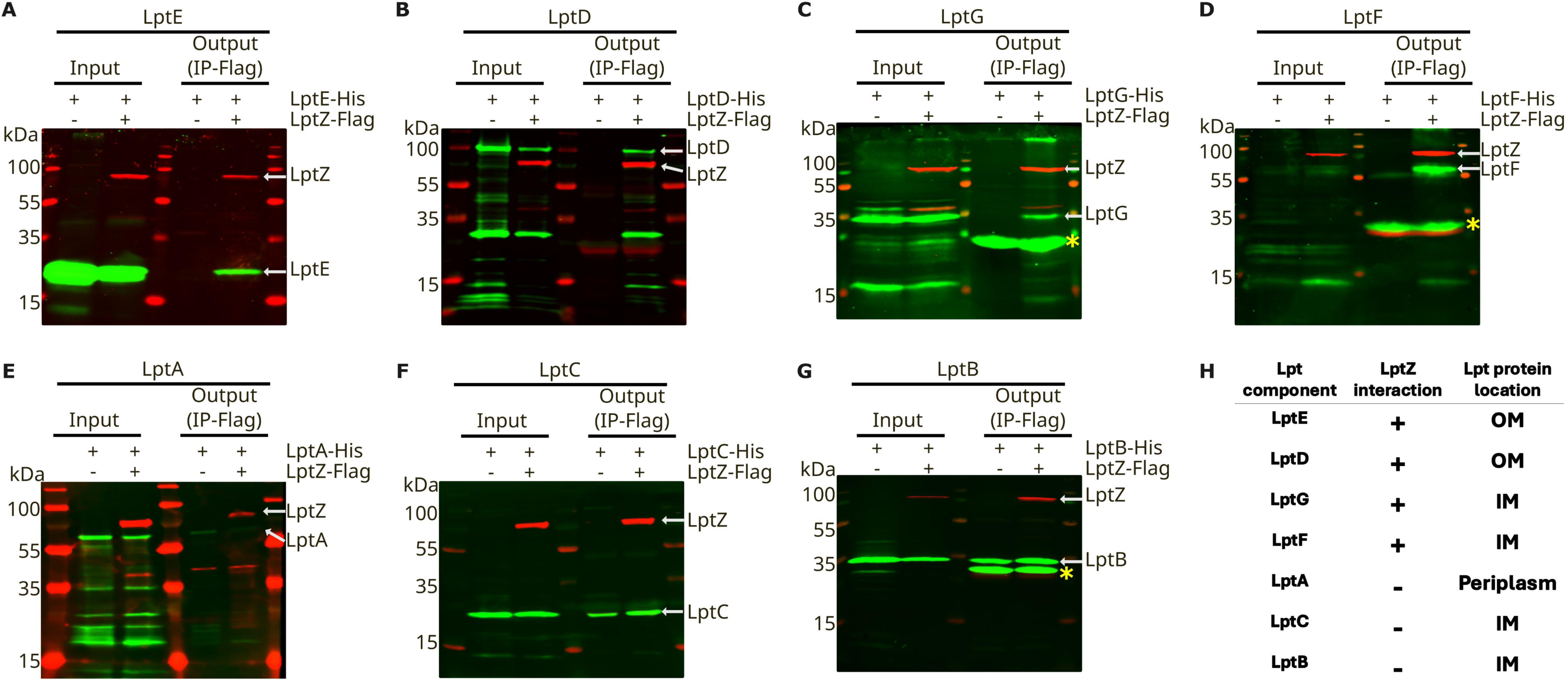
LptZ directly interacts with LptD, LptE, LptF and LptG. **(A-G)** LptZ selectively interacts with components of the Lpt transenvelope complex. **(A-G)** His-tagged LptA-G were co-expressed in *E. coli* with either empty vector (pBAD33.1) or FLAG-tagged LptZ (pBAD33.1:LptZ-FLAG). Lysates were subjected to anti-FLAG pulldown and immunoblotted with anti-His and anti-FLAG antibodies. Enrichment of His-tagged LptE (A), LptD (B), LptG (C), and LptF (D) in the presence of LptZ, compared to empty vector, indicates specific interactions. No enrichment of LptA (E), LptC (F), or LptB (G) was observed in the presence of LptZ compared to the empty vector, suggesting the lack of interaction with LptZ. Yellow stars indicate nonspecific bands from the anti-FLAG resin, which are absent in the input lanes. **(H)** Summary of the Co-IP analysis showing the Lpt proteins that co-purified with LptZ (+) and those for which no interaction was detected (-). Experiments were independently repeated at least three times with similar results; one representative result is shown.

**Figure S6.**
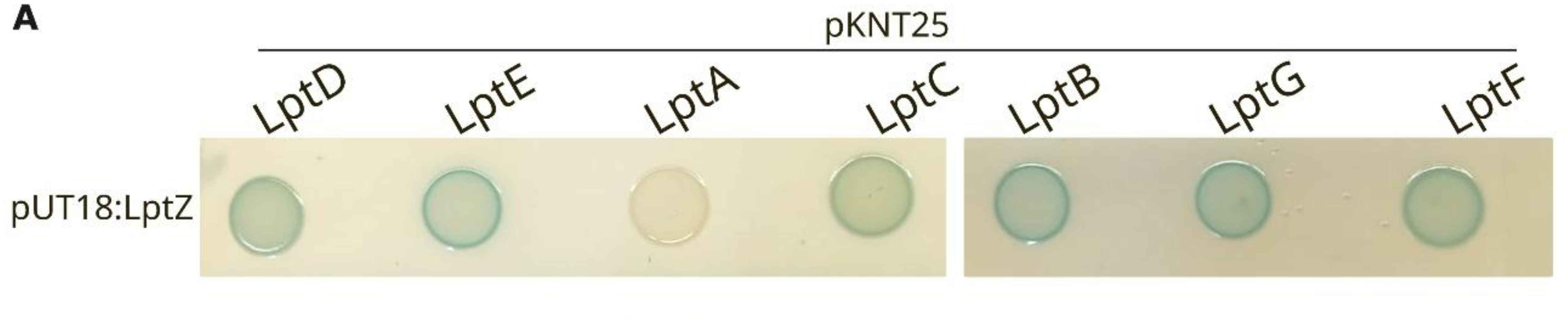
Bacterial two-hybrid assay to assess protein-protein interactions. Putative candidate proteins were fused to the T25 domain, and LptZ was fused to the T18 domain of *Bordetella pertussis* adenylate cyclase. Plasmids encoding the T18- and T25-fusion constructs were co-transformed into the *E. coli* BTH101 reporter strain. Protein-protein interactions were detected by spotting transformants onto LB agar plates supplemented with X-Gal and IPTG, and blue colonies indicate positive protein-protein interactions. Experiments were independently repeated at least three times with similar results; one representative result is shown.

**Figure S7.**
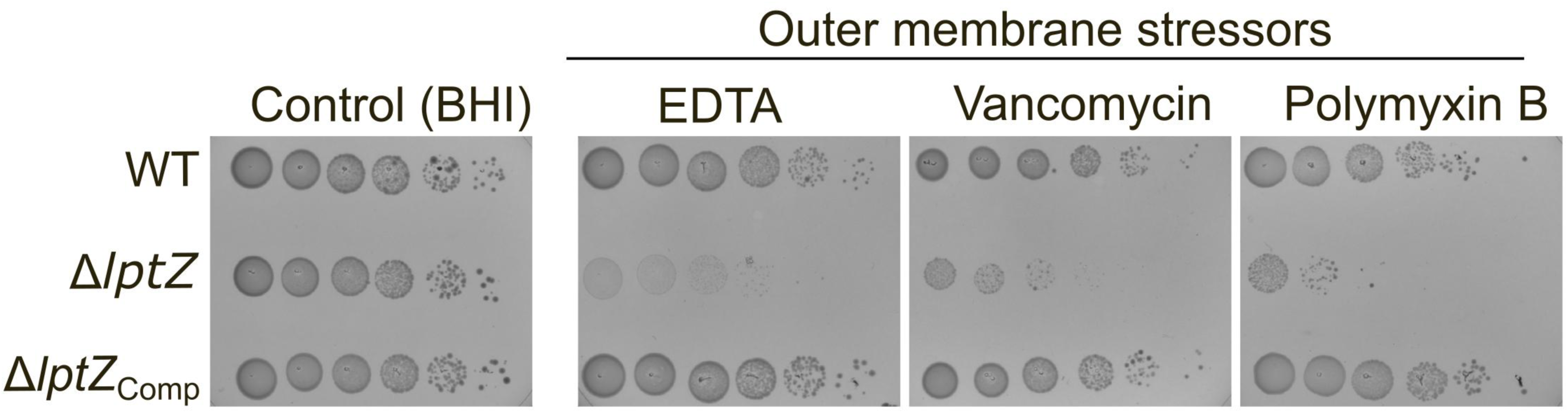
LptZ is required for outer-membrane integrity and resistance to envelope stress: Serial 10-fold dilutions of WT, Δ*lptZ* mutant, and complemented *Bt* strains were spotted on BHI agar with 1 mM EDTA, 16 µg/mL vancomycin, or 10 µg/mL polymyxin B. BHI alone served as a control. Plates were incubated anaerobically at 37 ⁰C for 48 h. Representative results from three independent experiments are shown here.

**Figure S8:**
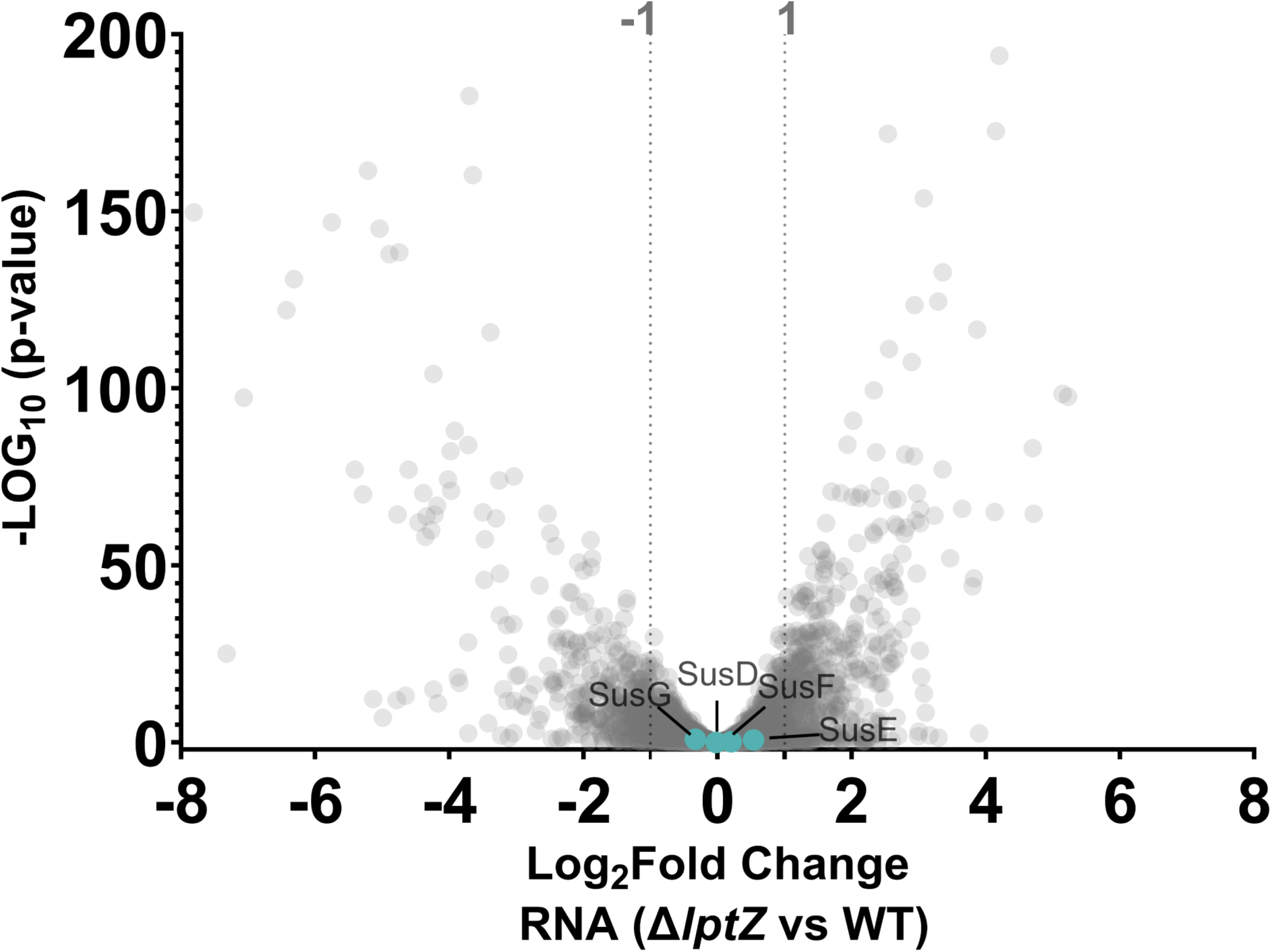
Volcano plot comparing transcript levels between WT and Δ*lptZ*. Dotted lines at log₂(fold change) ±1 indicate unchanged expression. All *sus* genes fall within this range, showing no transcriptional change in Δ*lptZ*.

**Figure S9.**
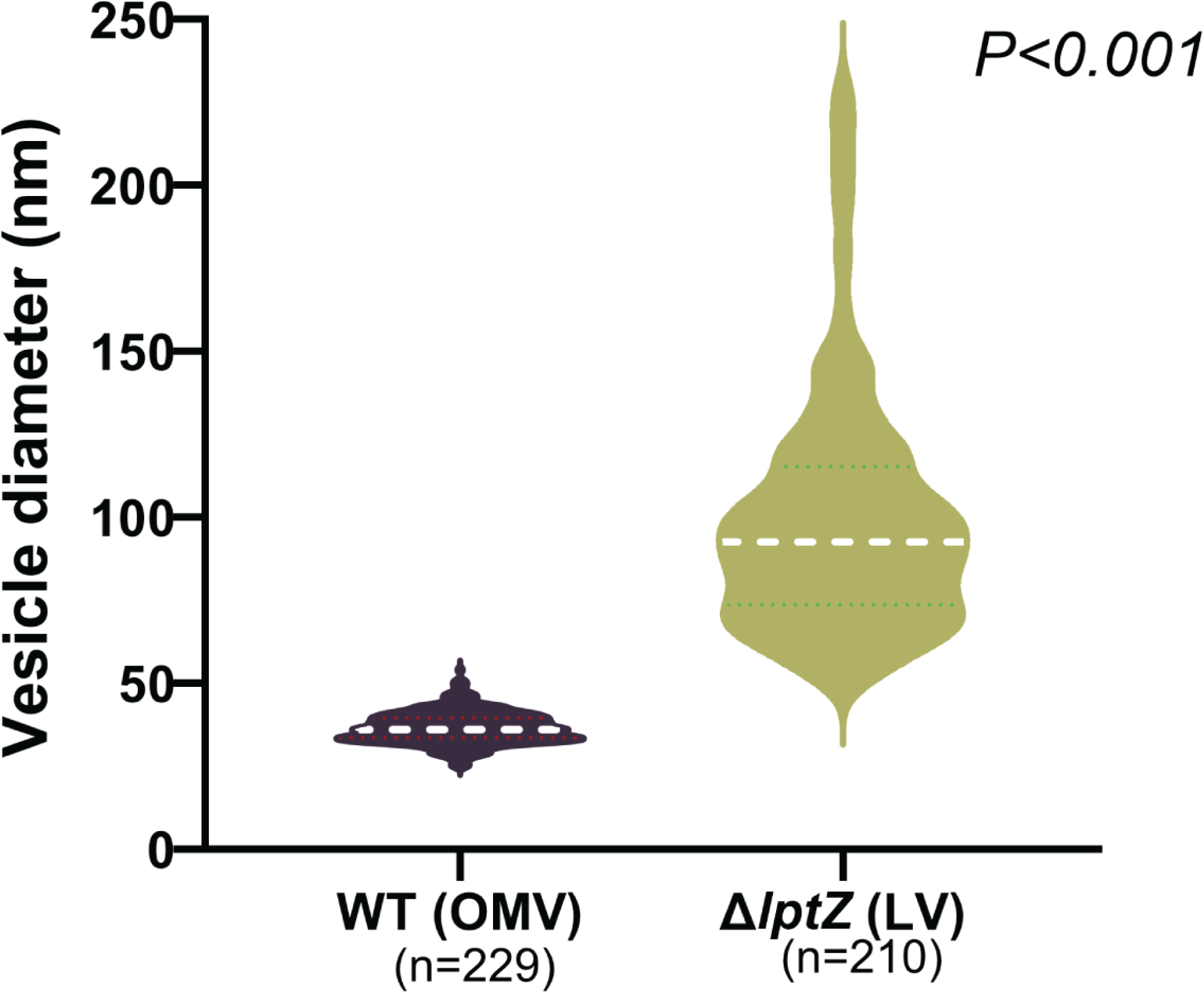
Vesicle diameters measured from cryo-ET images show a uniform size distribution in WT-derived OMVs, whereas LVs from the Δ*lptZ* mutant are highly heterogeneous and larger in size.

**Figure S10:**
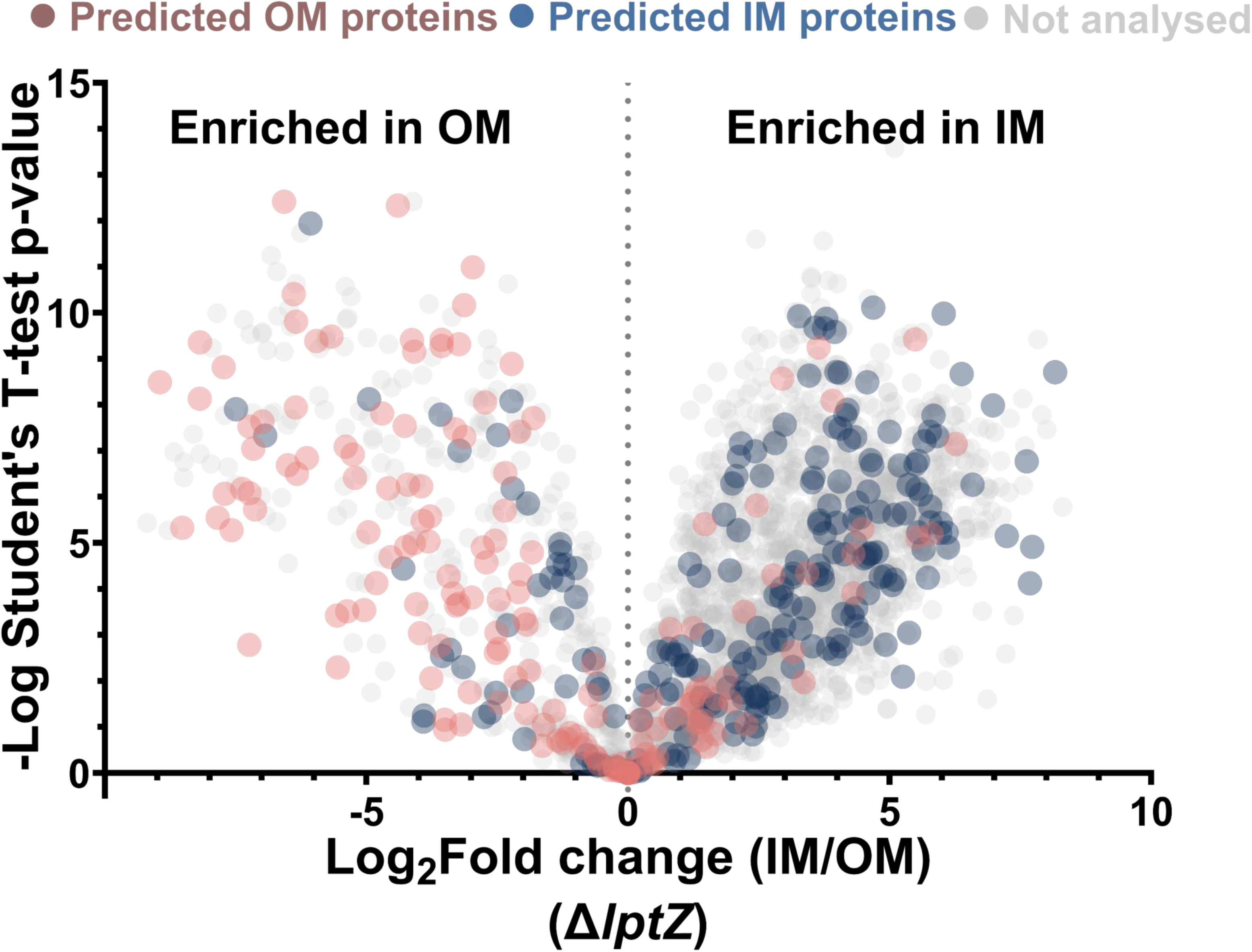
IM and OM separation in ΔLptZ. Volcano plot of isolated IM and OM proteomes of ΔLptZ, highlighting enrichment of predicted IM-specific proteins (blue circle) and OM markers (maroon circle) in their respective fractions.

**Figure S11:**
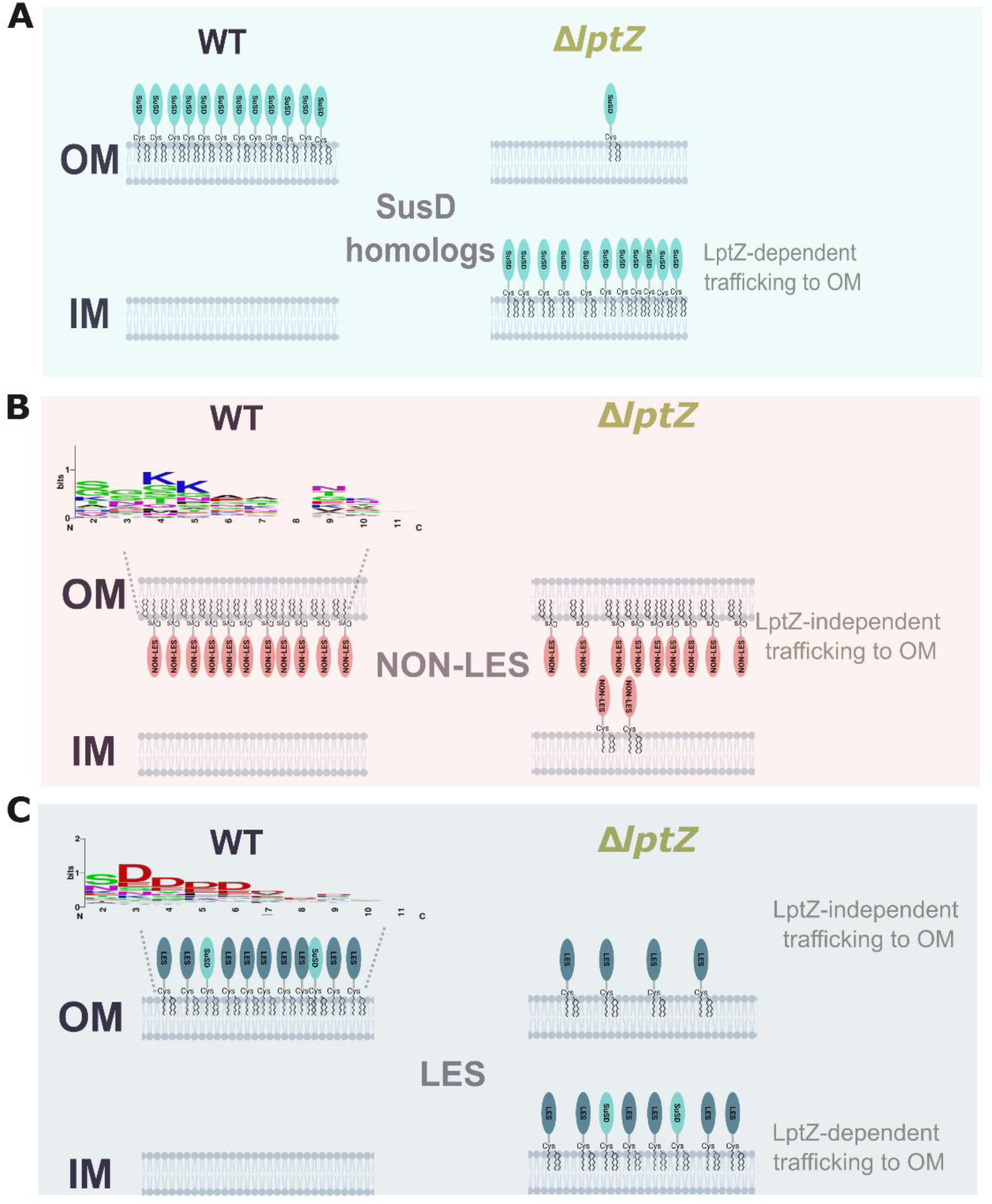
Schematic representation of lipoprotein transport mechanism, summarized from Figure 5. **(A)** In the WT strain, all SusD homologs localize to the outer surface of the OM, whereas in the Δ*lptZ* mutant, most SusD homologs are absent from the OM and retained in the IM as shown in Figure 5B. **(B)** Non-LES lipoproteins are largely unaffected in the Δ*lptZ* mutant and continue to localize to the periplasmic side of the OM, as shown in Figure 5D. **(C)** LES-containing lipoproteins, including SusD homologs, typically localize to the cell surface. In Δ*lptZ*, a subset accumulates in the IM and is absent from the cell surface, indicating impaired trafficking. In contrast, others still reach the surface, suggesting LptZ-independent transport for this group as shown in Figure 5E. Sequence logos representing the ten residues immediately downstream of the conserved lipoprotein attachment cysteine are shown for each group.

**Figure S12:**
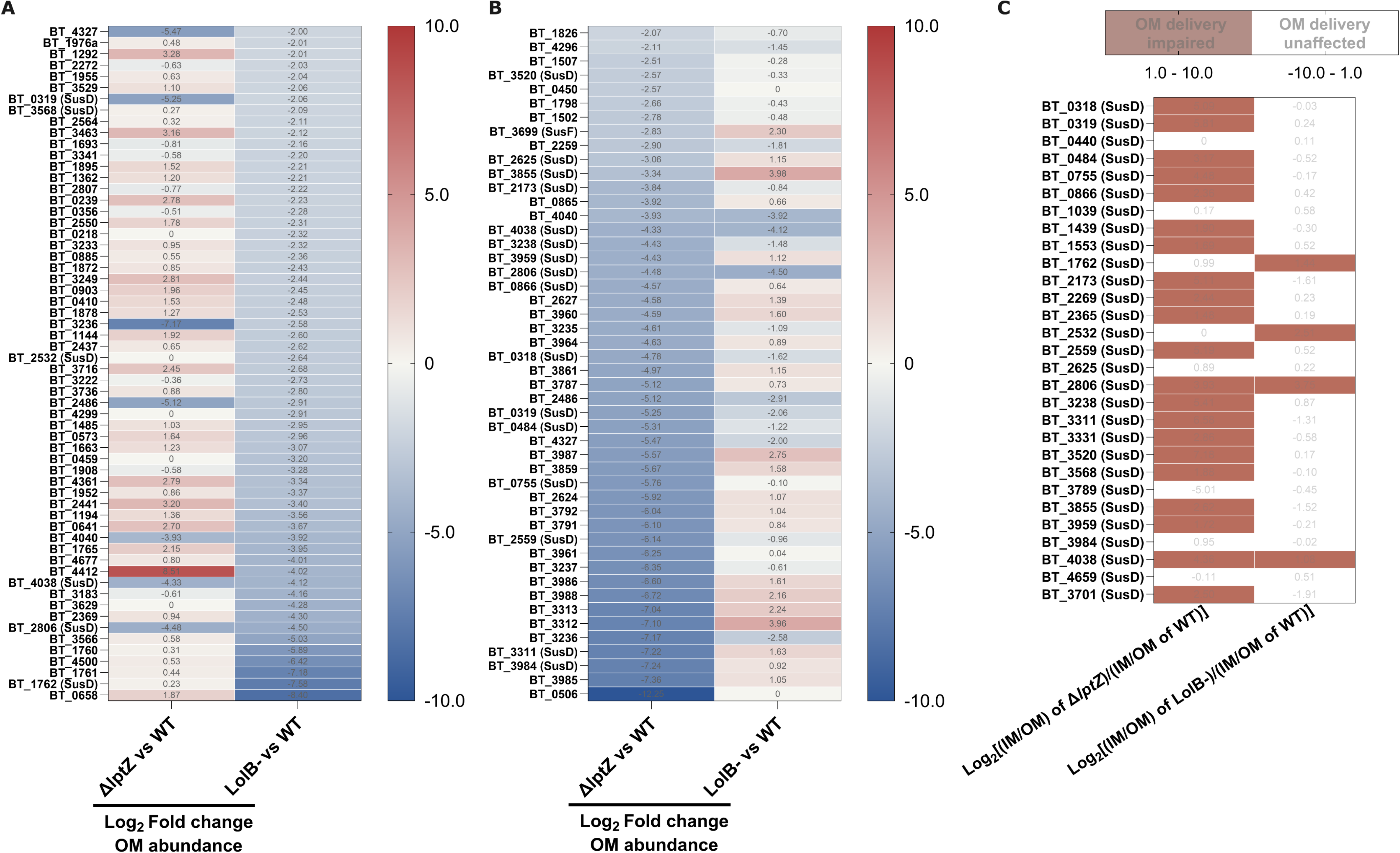
LptZ and LolB differentially affect OM lipoprotein trafficking. **(A–B)** Heat maps comparing changes in OM lipoprotein abundance in Δ*lptZ* and LolB- relative to WT. **(A)** Lipoproteins showing the strongest decreases in OM abundance in LolB- (log_2_ fold change < −2) compared to WT are shown, with their corresponding changes in Δ*lptZ*. **(B)** Conversely, lipoproteins most strongly depleted from the OM in LolB- (log₂ fold change < −2) are shown, together with their corresponding changes in Δ*lptZ*. The largely distinct patterns of lipoprotein depletion indicate that the effects of Δ*lptZ* are not attributable to a general disruption of lipoprotein trafficking and differ from those caused by loss of LolB. **(C)** Comparison of SusD-like lipoprotein trafficking defects in ΔlptZ and LolB-. Relative changes in subcellular distribution were calculated as log₂[(IM/OM ratio in mutant)/(IM/OM ratio in WT)] and plotted as a heat map. Positive values (>1) indicate increased IM retention and impaired OM delivery, whereas values (<1) indicate no substantial trafficking defect. Most SusD-like lipoproteins show increased IM/OM ratios in Δ*lptZ* but not in LolB-, consistent with a specific requirement for LptZ in SusD-like lipoprotein delivery to the OM.

**Figure S13:**
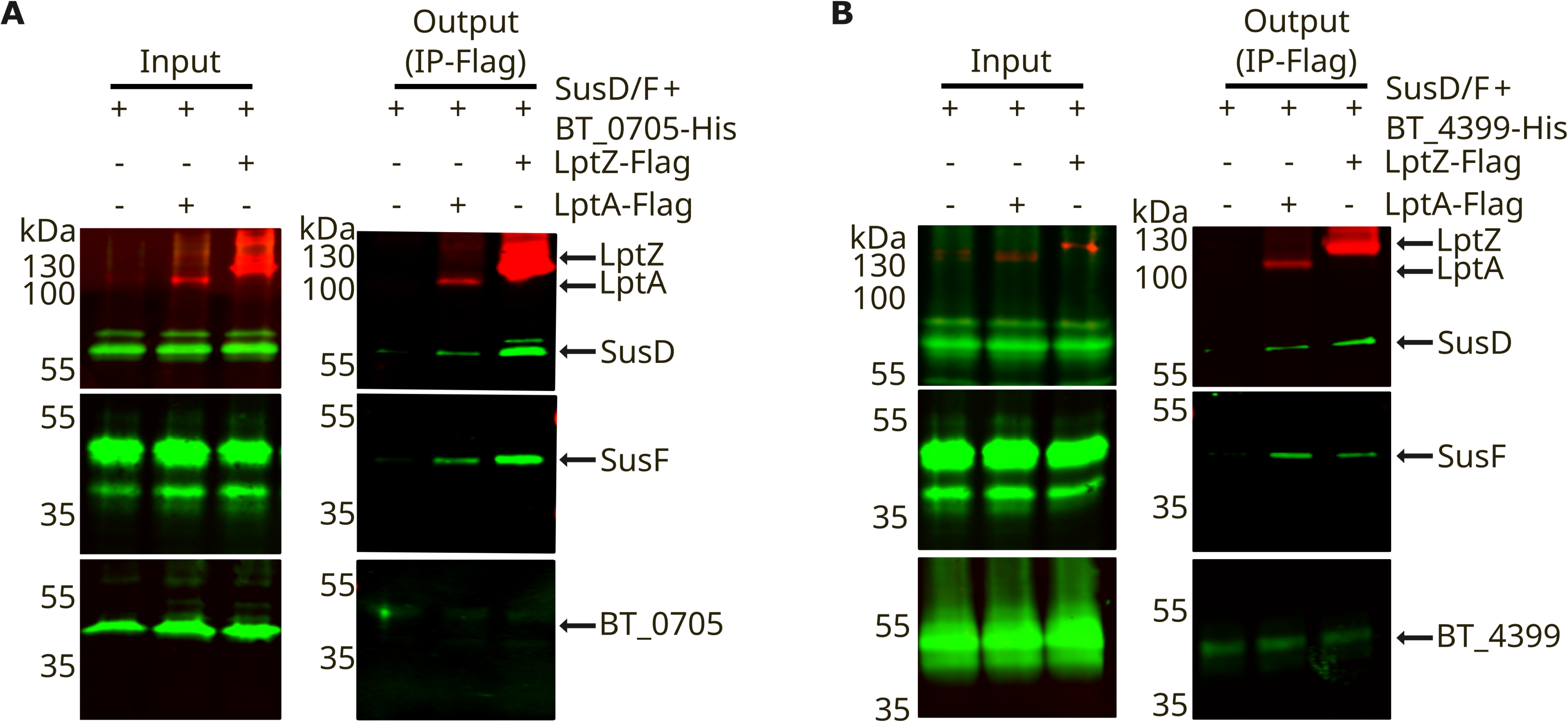
LES-containing lipoproteins selectively associate with the Lpt machinery. LptA and LptZ, each carrying a genomically encoded C-terminal 2xFLAG tag, were used as baits. **(A-B)** Maltose-induced SusD and SusF served as LES-containing substrates, whereas BT_0705 and BT_4399 represented non-LES outer-membrane lipoproteins and were constitutively expressed with C-terminal His tags. SusD and SusF were monitored with protein-specific antibodies, and BT_0705 and BT_4399 with an anti-His antibody. Both SusD and SusF were co-purified after DSP-mediated crosslinking with LptA and LptZ, whereas BT_0705 and BT_4399 were not relative to the untagged WT control. Experiments were independently repeated at least three times with similar results; one representative result is shown.

**Figure S14:**
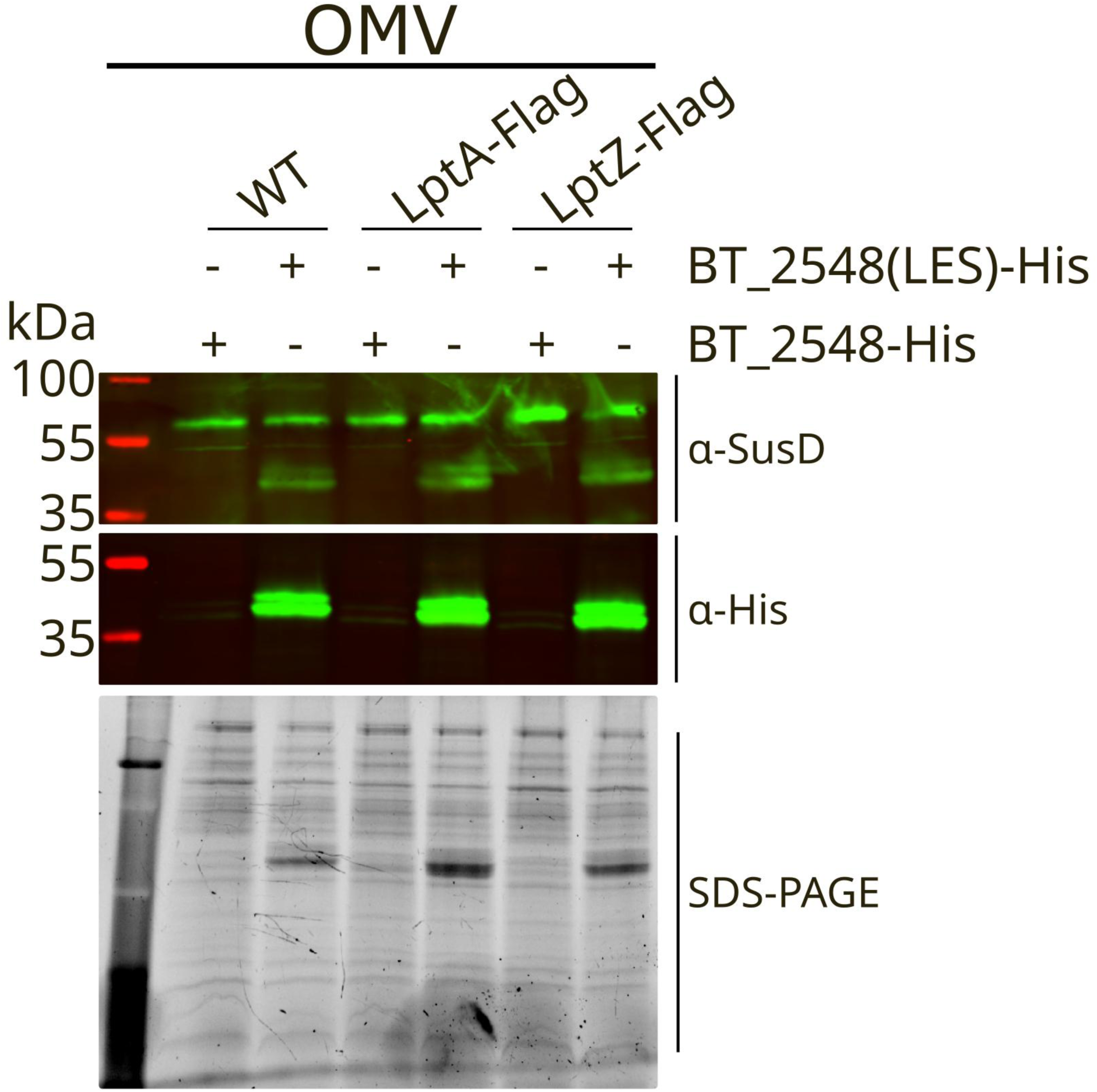
The SusD LES motif promotes incorporation of BT_2548 into OMVs. The native N-terminal non-LES sequence of BT_2548 was replaced with the LES sequence from SusD to generate BT_2548(LES). Both BT_2548 variants were C-terminally His-tagged. The resulting construct was introduced into WT and strains carrying genomically encoded C-terminal 2xFLAG tags on LptA or LptZ, and OMVs were isolated. **Upper panel:** Western blotting with an anti-SusD antibody confirmed comparable SusD abundance across strains. **Middle panel:** Immunoblotting with an anti-His antibody showed that replacement of the native non-LES with the SusD LES motif substantially increased the abundance of BT_2548 in the OMV fraction. **Lower panel:** SDS-PAGE analysis of the OMV fractions confirmed comparable OMV recovery across the strains. Experiments were independently performed at least twice with similar results; representative data are shown.

**Figure S15:**
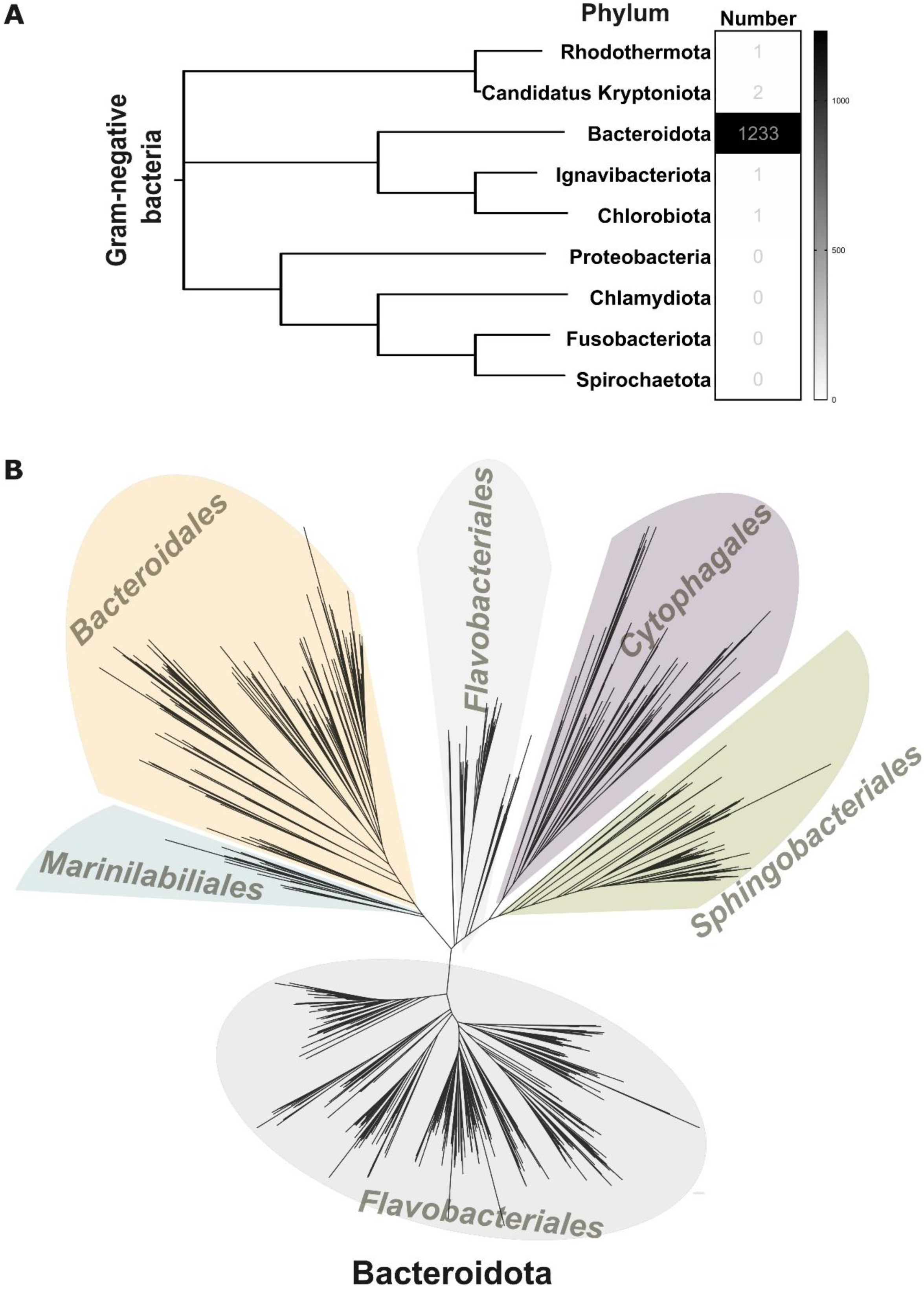
Functional conservation and phylogenetic distribution of LptZ. (**A)** Distribution of LptZ homologs across major phyla of Gram-negative bacteria. Numbers represent the count of LptZ homologs identified in each phylum during our analysis. **(B)** Phylogenetic tree of bacterial species encoding LptZ homologs. Branches are color-shaded by taxonomic order, including *Bacteroidales*, *Flavobacteriales*, *Cytophagales*, *Sphingobacteriales*, and *Marinilabiliales*.

**Table S1:** Top 50 abundant proteins in the OMV fraction of ΔLptZ versus WT.

| **Gene names** | **Protein names** | **Predicted cellular localization** | **Log<sub>2</sub> Fold Change OMV (ΔLptZ vs WT)** |
| --- | --- | --- | --- |
| BT_1998 | Anaerobic ribonucleoside-triphosphate reductase | Cytosol | 11.0921 |
| BT_3876 | 30S ribosomal protein S9 | Cytosol | 10.8605 |
| BT_0554 | Glutamine--fructose-6-phosphate aminotransferase | Cytosol | 10.8323 |
| BT_2829 | Arginine-tRNA ligase | Cytosol | 10.7537 |
| BT_4314 | 50S ribosomal protein L21 | Cytosol | 10.7233 |
| BT_2713 | 30S ribosomal protein S8 | Cytosol | 10.475 |
| BT_1499 | DNA-binding protein HU | Cytosol | 10.4695 |
| BT_3877 | 30S ribosomal protein S2 | Cytosol | 10.4376 |
| BT_3430 | 30S ribosomal protein S20 | Cytosol | 10.2642 |
| BT_3995 | Alanine--tRNA ligase | Cytosol | 10.1802 |
| BT_2714 | 30S ribosomal protein S14 | Cytosol | 10.157 |
| BT_0578 | Excinuclease ABC subunit A | Cytosol | 10.1304 |
| BT_3872 | Pseudouridine synthase | Cytosol | 10.0946 |
| BT_4101 | Alanine racemase | Cytosol | 9.93964 |
| BT_0425 | 50S ribosomal protein L20 | Cytosol | 9.89111 |
| BT_1829 | Chaperonin GroEL | Cytosol | 9.87164 |
| BT_3611 | Glycine--tRNA ligase | Cytosol | 9.8405 |
| BT_2724 | 50S ribosomal protein L2 | Cytosol | 9.77836 |
| BT_2710 | 30S ribosomal protein S5 | Cytosol | 9.75901 |
| BT_1535 | ABC transporter ATP-binding protein | Inner membrane | 9.74155 |
| BT_2701 | DNA-directed RNA polymerase subunit alpha | Cytosol | 9.68099 |
| BT_4610 | Protein RecA (Recombinase A) | Cytosol | 9.61258 |
| BT_1634 | Putative aminoglycoside phosphotransferase | Not determined | 9.51376 |
| BT_2715 | 50S ribosomal protein L5 | Cytosol | 9.50339 |
| BT_0408 | YtxH domain-containing protein | membrane | 9.4828 |
| BT_0565 | Putative heat shock protein |  | 9.31365 |
| BT_4362 | Protein translocase subunit SecA | Inner membrane | 9.30967 |
| BT_0871 | Diacylglycerol kinase-like, catalytic region | Cytosol | 9.29115 |
| BT_2416 | GTP cyclohydrolase-2 | Cytosol | 9.24956 |
| BT_0694 | ABC transporter, ATP-binding protein | Inner membrane | 9.24184 |
| BT_4002 | ATPase, ParA family | Cytosol | 9.23574 |
| BT_2703 | 30S ribosomal protein S11 | Cytosol | 9.23177 |
| BT_4041 | DUF349 domain-containing protein | Cytosol | 9.21363 |
| BT_2711 | 50S ribosomal protein L18 | Cytosol | 9.19412 |
| BT_2552 | Probable butyrate kinase (BK) | Cytosol | 9.17372 |
| BT_2721 | 30S ribosomal protein S3 | Cytosol | 9.13026 |
| BT_1747 | Pyruvate-flavodoxin oxidoreductase | Cytosol | 9.095 |
| BT_2720 | 50S ribosomal protein L16 | Cytosol | 9.03584 |
| BT_0623 | UDP-glucose 4-epimerase | Cytosol | 9.00702 |
| BT_0837 | ATP-dependent protease | Cytosol | 8.99484 |
| BT_3214 | S-adenosylmethionine:tRNA<br>ribosyltransferase-isomerase | Cytosol | 8.98287 |
| BT_2717 | 50S ribosomal protein L14 | Cytosol | 8.97322 |
| BT_3843 | ATP-dependent Clp protease ATP-binding<br>subunit ClpX | Cytosol | 8.85366 |
| BT_2704 | 30S ribosomal protein S13 | Cytosol | 8.80135 |
| BT_3054 | Succinate dehydrogenase flavoprotein subunit | Inner<br>membrane | 8.77293 |
| BT_0557 | carbamoyl-phosphate synthase | Cytosol | 8.76847 |
| BT_2734 | DNA-directed RNA polymerase subunit beta | Cytosol | 8.7472 |
| BT_2787 | 50S ribosomal subunit assembly factor BipA | Cytosol | 8.68427 |
| BT_4372 | Probable dual-specificity RNA<br>methyltransferase RlmN | Cytosol | 8.5796 |
| BT_2162 | 30S ribosomal protein S18 | Cytosol | 8.53864 |

**Table S2:** Proteins co-purified with LptZ and LptA in the presence or absence of the crosslinker DSP, as shown in Fig. 3C-E. Lpt machinery components are indicated in green, LptZ in blue, and proteins co-purified with both LptA and LptZ in the presence of DSP in pink.

|  | Bait proteins (Conditions) |  |  | Predicted function |
| --- | --- | --- | --- | --- |
|  | LptZ (- DSP) | LptZ (+DSP) | LptA (+DSP) |  |
| **LptD**<br>(BT_2133) | ✓ | ✓ | ✓ | **LPS transport** |
| **LptE**<br>(BT_4376) | ✓ | ✓ | ✓ | **LPS transport** |
| **LptA**<br>(BT_3849) | ✓ | ✓ | ✓ | **LPS transport** |
| **LptZ**<br>(BT_3341) | ✓ | ✓ | ✓ | **Unknown** |
| **BT_0154** | X | ✓ | ✓ | **Unknown** |
| **BT_4000** | X | ✓ | ✓ | **Unknown** |
| **BT_4562** | X | ✓ | ✓ | **Unknown** |
| **BT_4563** | X | ✓ | ✓ | **Unknown** |
| **BT_0328** | X | ✓ | ✓ | **Unknown** |
| **BT_1085** | X | ✓ | ✓ | **Unknown** |
| BT_0586 | ✓ | X | X | Unknown |
| BT_0550 | ✓ | X | X | Unknown |
| BT_2708 | ✓ | X | X | Unknown |
| BT_1843 | ✓ | X | X | Unknown |
| BT_4493 | ✓ | X | X | Unknown |
| BT_1573 | X | ✓ | X | Unknown |
| BT_0319 | X | ✓ | X | Unknown |
| BT_2758 | X | ✓ | X | Unknown |
| **LptF**<br>**(BT_2417)** | **X** | **X** | **✓** | **LPS transport** |
| **LptG**<br>**(BT_0834)** | **X** | **X** | **✓** | **LPS transport** |
| **LptC**<br>**(BT_4369)** | **X** | **X** | **✓** | **LPS transport** |
| **LptB**<br>**(BT_3837)** | **X** | **X** | **✓** | **LPS transport** |
| BT_0809 | X | X | ✓ | Unknown |
| BT_2845 | X | X | ✓ | Unknown |
| BT_0747 | X | X | ✓ | Unknown |
| BT_3784 | X | X | ✓ | Unknown |
| BT_3999 | X | X | ✓ | Unknown |

**Table S3.**
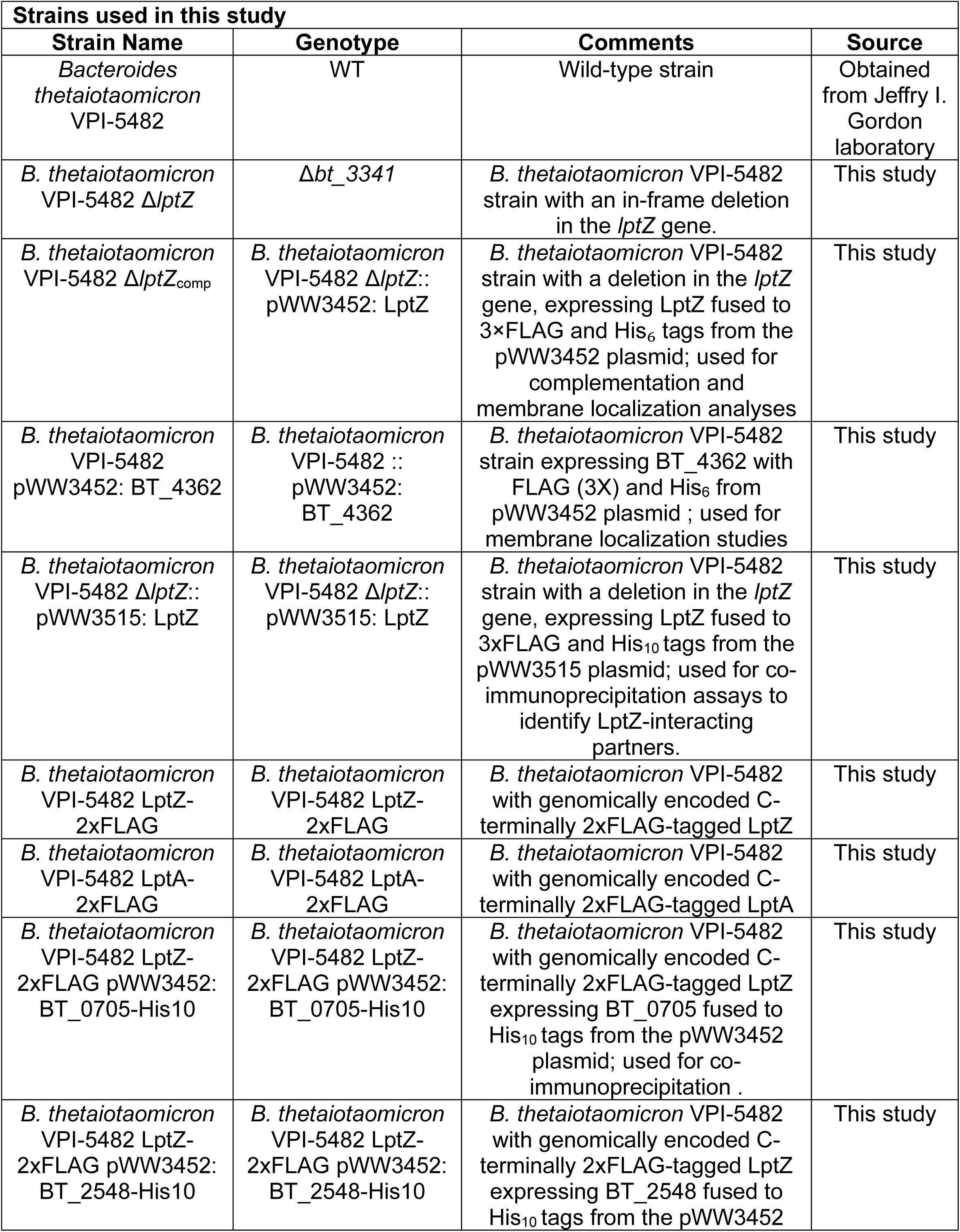
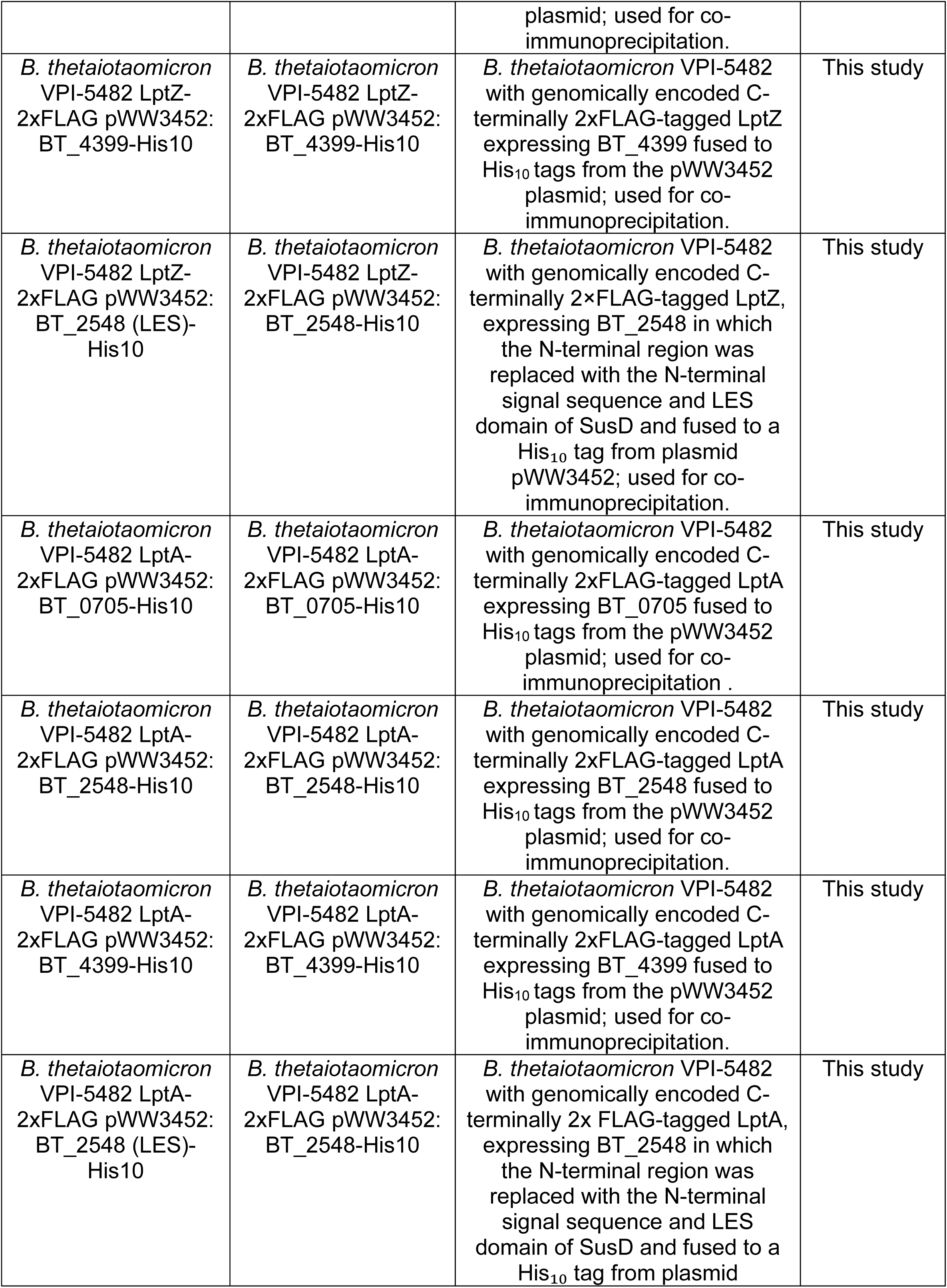
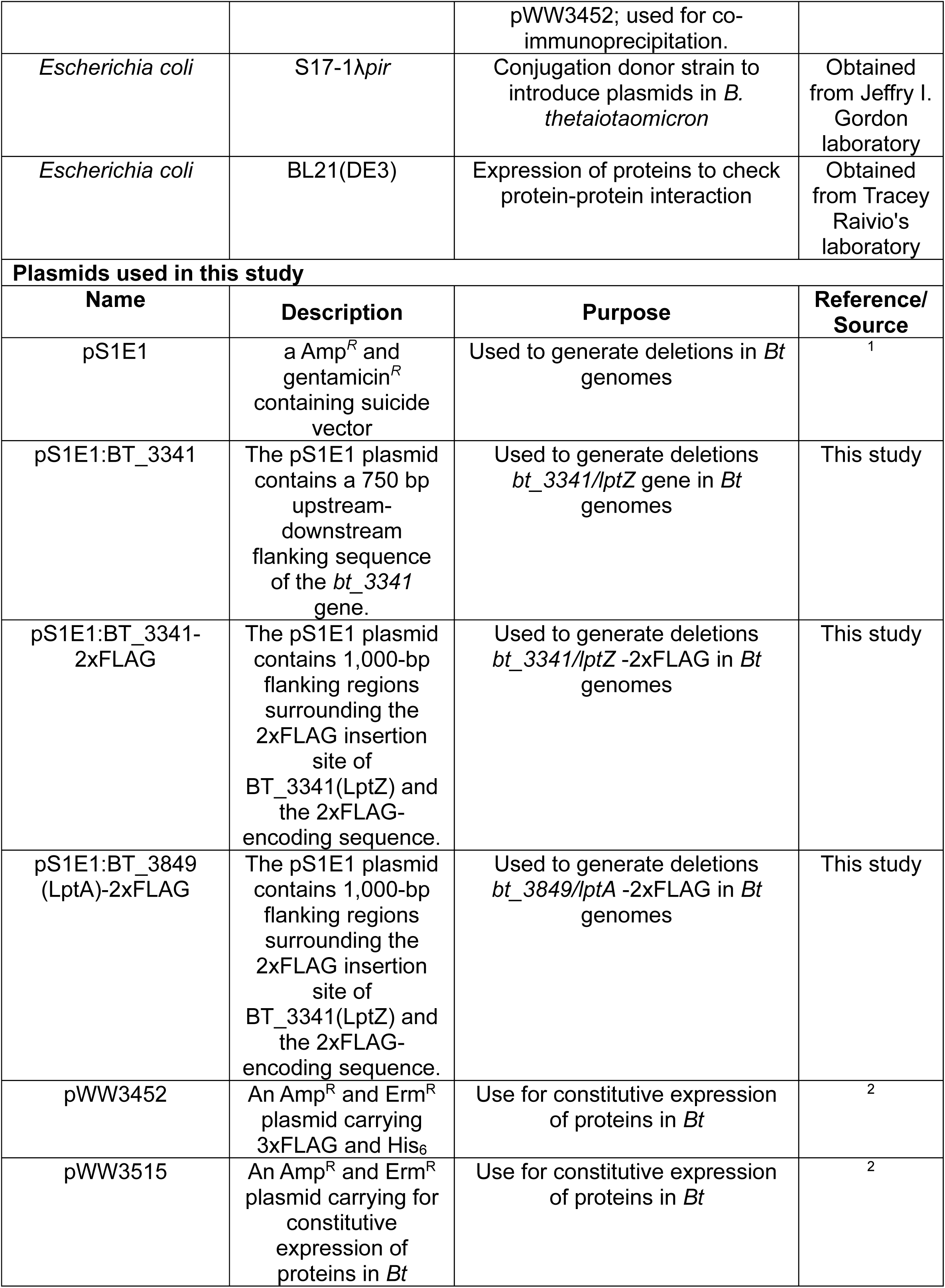
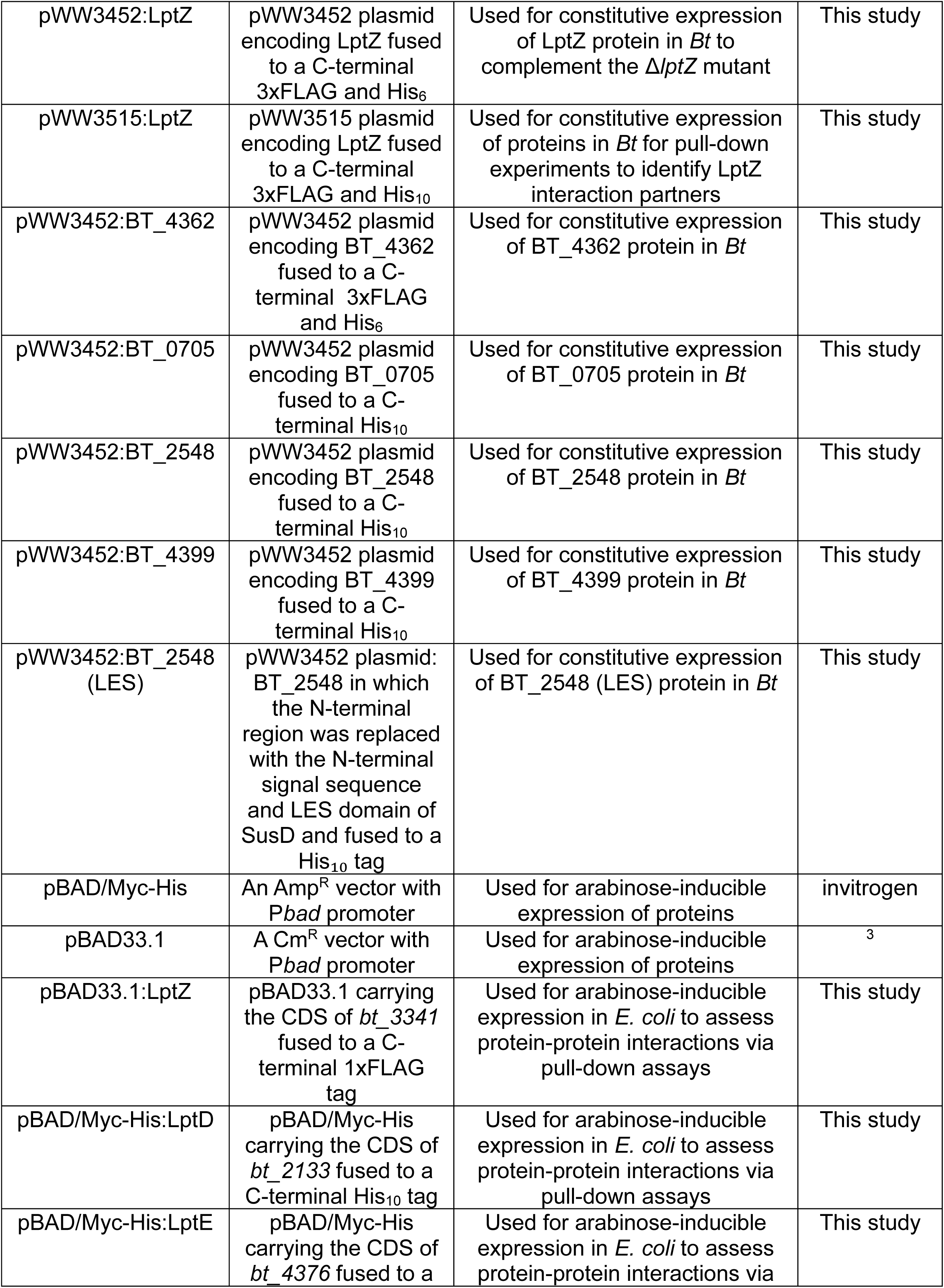
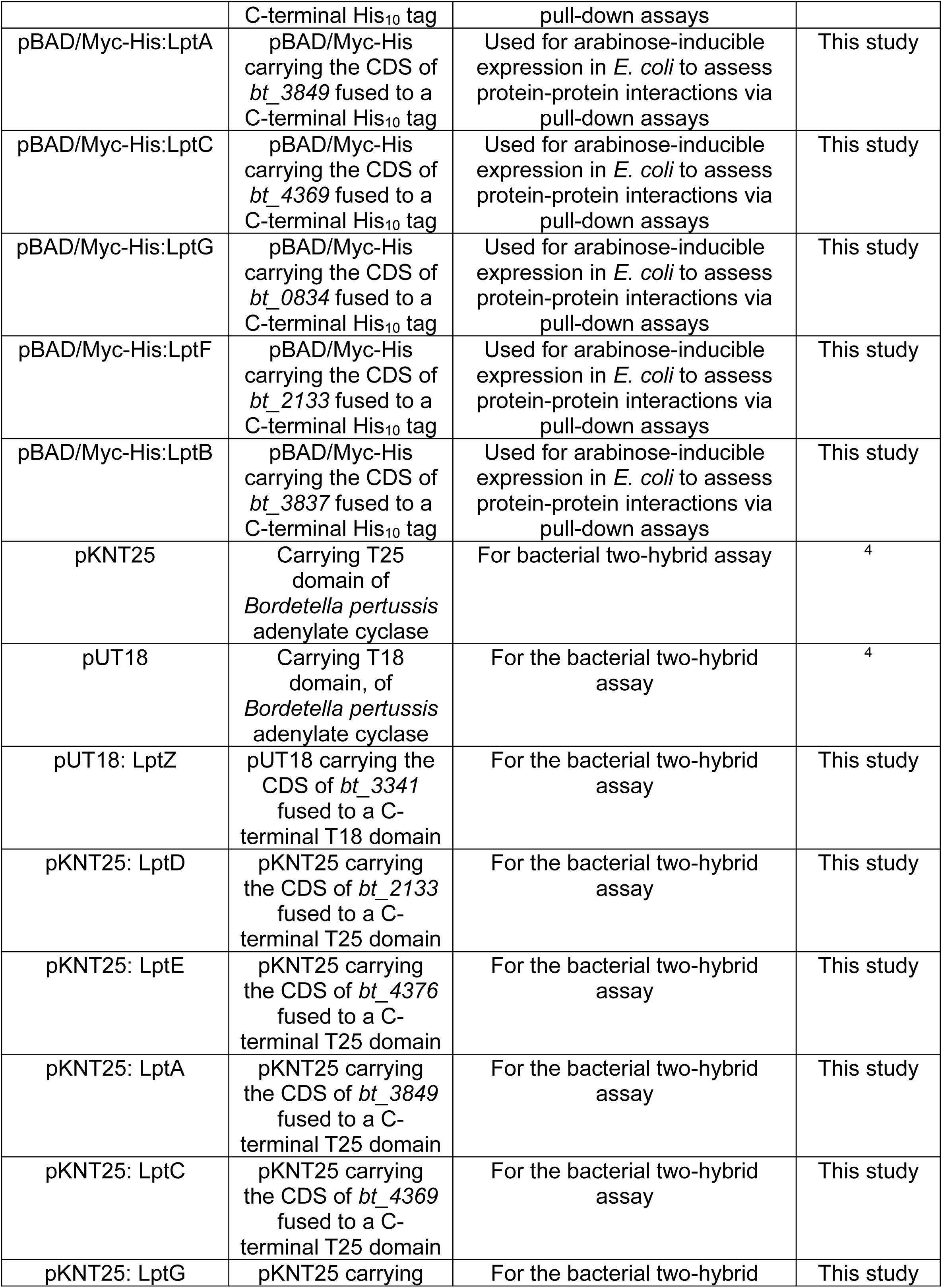
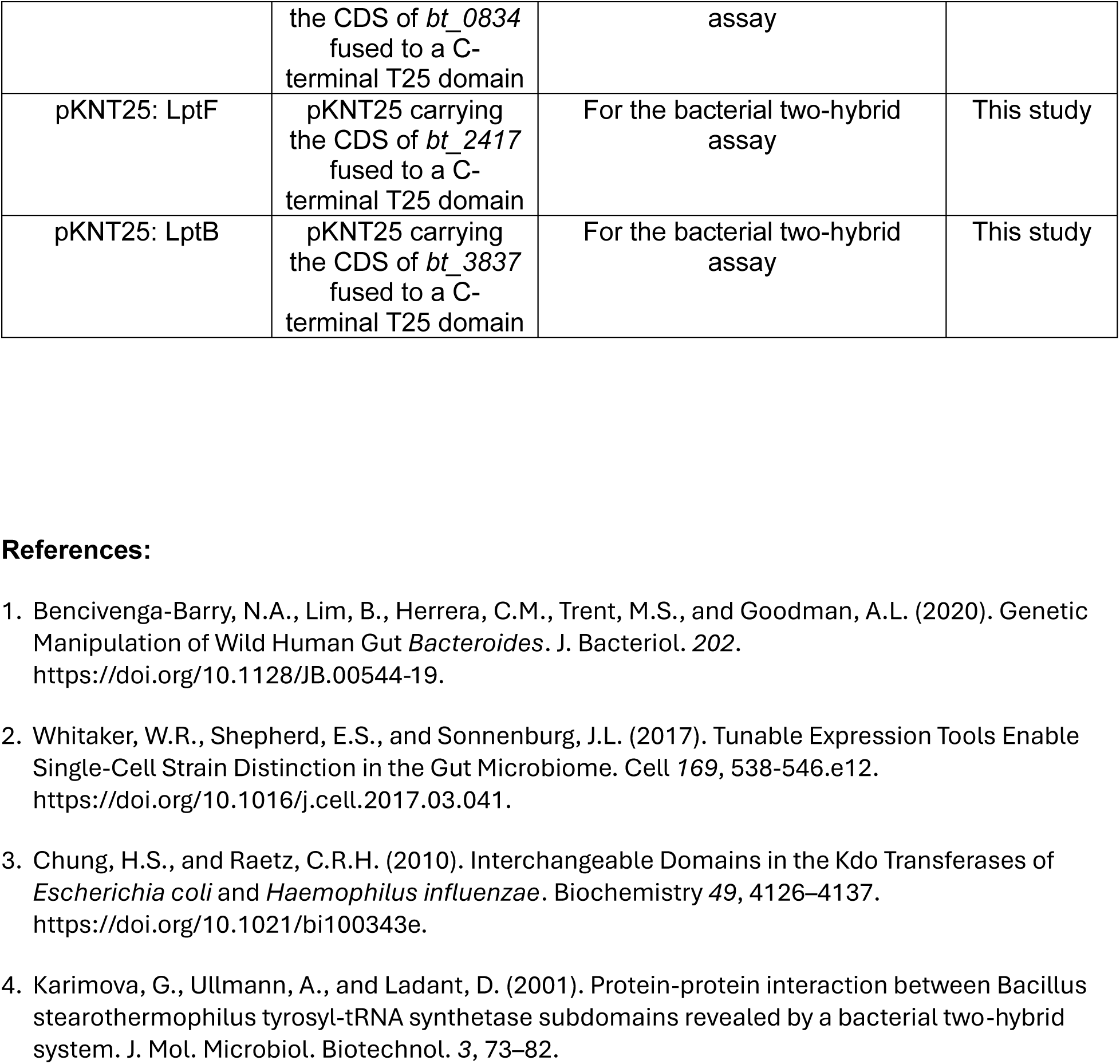
A list of bacterial strains and plasmids used in this work.

## Notes

### Competing Interest Statement

The authors have declared no competing interest.

